# Disrupting D2-NMDA receptor heteromerization blocks the rewarding effects of cocaine but preserves natural reward processing

**DOI:** 10.1101/2021.01.25.428078

**Authors:** Andry Andrianarivelo, Estefani Saint-Jour, Paula Pousinha, Sebastian P. Fernandez, Anna Petitbon, Veronique De Smedt-Peyrusse, Nicolas Heck, Vanesa Ortiz, Marie-Charlotte Allichon, Vincent Kappès, Sandrine Betuing, Roman Walle, Ying Zhu, Charlène Joséphine, Alexis-Pierre Bemelmans, Gustavo Turecki, Naguib Mechawar, Jonathan A Javitch, Jocelyne Caboche, Pierre Trifilieff, Jacques Barik, Peter Vanhoutte

**Affiliations:** CNRS, UMR 8246, Neuroscience Paris Seine, F-75005, Paris, France; INSERM, UMR-S 1130, Neuroscience Paris Seine, Institute of Biology Paris Seine, F-75005, Paris, France; Sorbonne Université, UPMC Université Paris 06, UM CR18, Neuroscience Paris Seine, F-75005, Paris, France; Université Côte d’Azur, Nice, France; Institut de Pharmacologie Moléculaire & Cellulaire, CNRS UMR7275, Valbonne, France; Université Bordeaux, INRAE, Bordeaux INP, NutriNeuro, 33000, Bordeaux, France; Division of Molecular Therapeutics, New York State Psychiatric Institute, New York, NY 10032, USA; Department of Psychiatry, Columbia University, New York, NY 10032, USA; Commissariat à l’Énergie Atomique et aux Énergies Alternatives (CEA), Département de la Recherche Fondamentale, Institut de biologie François Jacob, MIRCen, and CNRS UMR 9199, Université Paris-Sud, Université Paris-Saclay, Neurodegenerative Diseases Laboratory, Fontenay-aux-Roses, France; Douglas Mental Health University Institute, Department of Psychiatry, McGill University, Montreal, QC, Canada; Department of Pharmacology, Columbia University, New York, NY 10032, USA

## Abstract

Addictive drugs increase dopamine in the nucleus accumbens (NAc), where it persistently shapes excitatory glutamate transmission and hijacks natural reward processing. Herein, we provide evidence, from mice to human, that an underlying mechanism relies on drug-evoked heteromerization of glutamate NMDA receptors (NMDAR) with dopamine receptor 1 (D1R) or 2 (D2R). Using temporally-controlled inhibition of D1R-NMDAR heteromerization, we unraveled their selective implication in early developmental phases of cocaine-mediated synaptic, morphological and behavioral responses. In contrast, preventing D2R-NMDAR heteromerization blocked the persistence of these adaptations. Importantly, interfering with these heteromers spared natural reward processing. Strikingly, we established that D2R-NMDAR complexes exist in human samples and showed that, despite a decreased D2R protein expression in the NAc, psychostimulant-addicts display a higher proportion of D2R forming heteromers with NMDAR. These findings contribute to a better understanding of molecular mechanisms underlying addiction and uncover D2R-NMDAR heteromers as targets with potential therapeutic value.

## Introduction

Drug addiction is characterized by compulsive patterns of drug-seeking and drug-taking behavior in spite of detrimental consequences and a high rate of relapse after withdrawal. A hallmark of addictive drugs is their ability to increase dopamine concentration in discrete brain regions, which persistently shapes excitatory glutamate transmission within the reward circuit, thereby hijacking natural reward processing (*1*, *2*). This calls for a better understanding of the precise molecular events underlying the detrimental interplay between dopamine and glutamate signaling triggered by drugs of abuse.

The enduring behavioral alterations induced by protracted drug exposure are largely believed to result from persistent drug-evoked neuronal adaptations within the striatum, especially in its ventral part, the nucleus accumbens (NAc) (*1*, *3*). The striatum is indeed a key target structure of drugs of abuse that integrates convergent glutamate inputs from limbic, thalamic and cortical regions, encoding components of drug-associated stimuli and environment, and dopamine signals that mediate reward prediction error and incentive values (*4*). Integration of dopamine and glutamate signals is achieved by the two segregated subpopulations of GABAergic medium-sized spiny neurons (MSN) expressing either the dopamine receptor (DAR) type 1 (D1R) or type 2 (D2R), although a fraction of MSN in the NAc expresses both receptors (*5*). Cell-type-specific manipulations of neuronal activity showed that inhibiting and activating D1R-MSN respectively dampens and potentiates long-term drug-evoked responses, in line with their “pro-reward” action (*2*, *6*–*9*). By contrast, the majority of studies supports an inhibitory role of D2R-MSN activation on drug-mediated adaptations (*6*, *7*, *10*–*12*). These studies, based on direct manipulations of MSN activity, were extremely instrumental to highlight the role of MSNs as putative players in drug-related behavioral adaptations. However, they do not establish how drugs of abuse persistently impact the functionality of each MSN subpopulation or the underlying cellular and molecular mechanisms. In this context, increasing evidence suggests that such a central role of MSN subpopulations in drug-induced behavioral adaptations originates, at least in part, from dopamine-dependent long-lasting changes at excitatory striatal synapses. Indeed, long-term potentiation of specific glutamatergic afferences impinging onto D1R-MSN induced by dopamine is responsible for both the induction and maintenance of long-lasting behavioral adaptations to repeated cocaine exposure (*13*–*15*). Interestingly, glutamate transmission onto D2R-MSN seems to be spared by cocaine exposure, but selectively altered during cocaine craving after long access to high doses of cocaine (*16*).

It is therefore timely to identify molecular mechanisms by which drug-evoked increases in dopamine can permanently hijack glutamate transmission onto MSN. Although a number of studies have described the crosstalk between D1R and glutamate receptor of the NMDA (NMDAR) subtype as a key player in the behavioral effects of psychostimulants (*2*, *17*–*21*, *21*, *22*), the underlying molecular mechanisms remain elusive. Moreover, the processes by which dopamine impairs D2R-MSN activity to promote long-lasting drug-induced reinforcement is yet unknown.

Heteromeric complexes formed between dopamine receptors (DAR) and glutamate NMDAR have been proposed as integrators of dopamine and glutamate signals in both MSN populations (*23*). Receptor heteromers are of particular interest, not only because of their ability to dynamically modulate the component receptor’s functions in time and space, but also because they exhibit functional properties distinct from the component receptors, making them attractive targets for the development of more selective pharmacological strategies (*24*–*27*). Most evidence generated to date regarding dopamine-NMDA receptor heteromer functions come from *in vitro* and *ex vivo* studies and their potential role in long-term drug-induced adaptations has been overlooked. D1R form heteromers with GluN1 subunits of NMDAR *in vitro* (*28*, *29*) and *in vivo* in the striatum (*30*, *31*) and have been shown to allow the facilitation of NMDAR signaling by dopamine in D1R-MSN *ex vitro* on striatal slices (*31*). By contrast, the binding of D2R to GluN2B subunits of NMDAR mediates the inhibition of NMDA currents by dopamine in D2R-MSN and controls acute stereotypic locomotor responses to high cocaine doses (*32*). DAR-NMDAR heteromers therefore appear as putative molecular platforms that mediate crosstalk between dopamine and glutamate transmission onto MSNs. However, whether such receptor heteromers might constitute molecular substrates by which drugs of abuse enduringly alter glutamate transmission and trigger long-lasting behavioral alterations has not been studied.

We therefore investigated D1R-GluN1 and D2R-GluN2B heteromerization in the striatum *in vivo* in response to repeated cocaine exposure. We found that cocaine triggers a transient increase of D1R-GluN1 heteromerization in the entire striatum, which returns to baseline level upon withdrawal from the drug. By contrast, cocaine induces a stable heteromerization of D2R-Glu2NB that is mostly restricted to the NAc and persists over a withdrawal period. Using a temporally-controlled disruption these receptor heteromers, combined with electrophysiological recordings, imaging and behavioral assessments, we showed that D1R-GluN1 heteromerization controls the development of cocaine-evoked long-term synaptic plasticity and morphological changes in D1R-MSNs, as well as behavioral adaptations. In contrast, D2R-GluN2B heteromerization mediates the persistence of these adaptations after a withdrawal period followed by a re-exposure to the drug. Importantly, the targeting of either type of heteromers preserves natural reward processing. Strikingly, we found that such receptor complexes also exist in human post-mortem brain samples and showed that, despite a substantial decrease of D2R protein expression, psychostimulant-addicts display a significantly higher proportion of D2R that form heteromers with GluN2B in the NAc. Our results support a model by which the heteromerization of dopamine and glutamate receptors induced by drugs of abuse in D1R- and D2R-MSNs is a key endogenous molecular event underlying the detrimental interplay between these two neurotransmitter systems in drug addiction. The role of D2R-GluN2B heteromers in the persistence of the sensitizing and rewarding effects of drugs makes them potential targets not only for addiction in humans, but also more broadly in multiple neuropsychiatric disorders.

## Results

### Behavioral sensitization to cocaine is associated with transient D1R-NMDAR heteromerization and prolonged D2R-GluN2B heteromerization in the NAc

Before studying the role of DAR-NMDAR heteromers in cocaine-evoked adaptations, we investigated whether a cocaine regimen that triggers persistent behavioral adaptations can modulate the formation of these receptor complexes *in vivo* in the striatum. Mice were subjected to five daily injections of cocaine (15 mg/kg), which elicits a progressive locomotor sensitization (Fig. 1A) that is known to persist for several weeks after cocaine withdrawal. This behavioral paradigm is a straightforward model to study the mechanisms involved in drug-induced behavioral adaptations (*2*, *33*). Mice were sacrificed one day after the last injection to detect endogenous DAR-NMDAR proximity in distinct striatal sub-regions through Proximity Ligation Assay (PLA) (*34*). The brightfield PLA assay yielded a brown punctate signal for D1R-GluN1 and D2R-GluN2B complexes that was absent when one of the two primary antibodies was omitted (Fig. 1B), similar to what was previously found when PLA was performed in DA receptor KO mice (*31*, *34*, *35*). Using this approach, we found that cocaine-treated mice displayed increased D1R-GluN1 heteromerization in the dorso-lateral (DL Str) and dorso-medial (DM Str) striatum, as well as in the nucleus accumbens core (NAc core) and shell (NAc shell) sub-divisions (Fig. 1C). Cocaine also increased D2R-GluN2B heteromerization, but primarily in the NAc, with a smaller effect in the DL Str (Fig. 1D). This increased heteromerization occurred in the absence of changes in global expression levels of the component receptors (Fig. S1A,B). Of note, and as previously observed (*36*), repeated cocaine exposure decreases expression levels of the synaptic scaffold protein PSD-95 (Fig. S1C),which could partly explain our results as the interaction of NMDAR and D1R with PSD-95, through partly overlapping domains, has been described to prevent D1R-GluN1 interaction (*37*).

**Figure 1.**
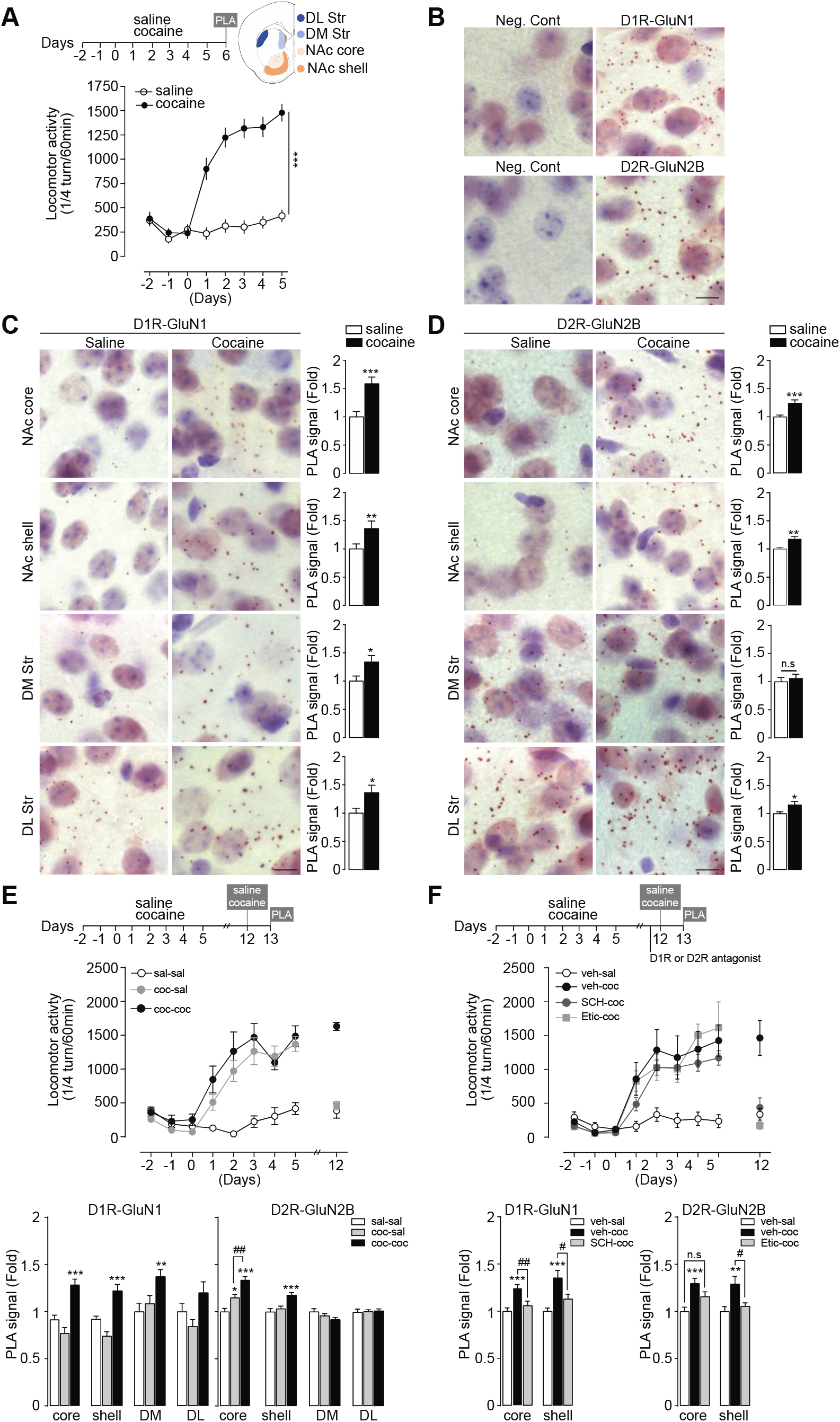
Behavioral sensitization to cocaine is associated with transient D1R-NMDAR heteromerization and prolonged D2R-GluN2B heteromerization in the NAc. (**A**) Experimental time frame (DL Str: dorso-lateral striatum; DMStr dorso-medial striatum; nucleus accumbens core(NAc core) and shell(NAc shell) andmeasurements of locomot or activity prior and during 5 days of saline or cocaine (15 mg/kg) injections. Two-way ANOVA, treatment effect, F (1, 24) = 67.79, *** P < 0.0001 saline vs cocaine on day 5, n=13 mice/group. (**B**) Example images of D1R-GluN1 and D2R-GluN2B heteromer PLA detection and related negative (Neg. Cont) showing the absence of signal when one primary antibody is omitted. (**C**) Detection and quantifications of D1R-GluN1 heteromerization in saline- and cocaine-treated groups. PLA signal is represented as fold increase normalized to the saline group. Two-sided Student’s *t*-test. * *P* < 0.05; ** *P* < 0.01; *** *P* < 0.001 saline vs cocaine, n=28-84 fields of view/structure (NAc core: 4, NAc shell: 12; DM: 6, DL: 6, fields of view/mice) from 7 mice/group. (**D**) Same as for (C) for D2R-GluN2B heteromerization. (**E**) Experimental time frame, measurements of locomotor activity and quantifications of D1R-GluN1 and D2R-GluN2Bheteromerization. PLA signal is represented as fold increase normalized to the saline group. One-way ANOVA. * *P* < 0.05;** *P* < 0.01;*** *P* < 0.001 saline-saline vs cocaine-saline/cocaine-cocaine, ## p<0.01 cocaine-saline vs cocaine-cocaine, n=28-84 fields of view/structure from 7 mice/group. (**F**) Same as for (E) except that mice received either a vehicle solution (veh) or the D1R antagonist SCH23390 (SCH) or the D2R antagonist Eticlopride (Etic) prior to a cocaine challenge. One-way ANOVA. * *P* < 0.05; ** *P* < 0.01; *** *P* < 0.001 vehicle-saline vs vehicle-cocaine, # p < 0.05, ## p<0.01 vehicle-cocaine vs SCH-cocaine or cocaine vs Etic-cocaine, n.s: not significant, n=28-84 fields of view/structure from 7 mice/group. (B-D) scale bar: 10 μm. Error bars denote s.e.m.

To study the kinetics of receptor heteromerization, mice were subjected to a cocaine-induced locomotor sensitization paradigm followed by one-week withdrawal and a challenge injection of saline or cocaine. We found that cocaine-mediated D1R-GluN1 heteromerization returned to baseline levels upon withdrawal but increased again, in all striatal sub-regions except in the DL-Str, after the challenge injection of cocaine (Fig. 1E).

Strikingly, D2R-GluN2B heteromerization appeared to be sustained over the withdrawal period specifically in the NAc core. The challenge injection of cocaine further increased D2R-Glu2NB heteromers in the NAc, but not in dorsal parts of the striatum (Fig. 1E). To assess the role of D1R or D2R stimulation for receptor heteromerization, mice underwent a cocaine-locomotor sensitization followed by a withdrawal. Before a cocaine challenge, mice received an intraperitoneal injection of D1R or D2R antagonist that blunted the expression of behavioral sensitization. This allowed us to show that the stimulation of D1R or D2R was mandatory for cocaine-induced D1R-GluN1 and D2R-GluN2B heteromerization, respectively (Fig. 1F). Altogether, these data show that behavioral sensitization to cocaine is associated with a dopamine receptor-dependent transient heteromerization of D1R-GluN1 in the whole striatum, whereas D2R-GluN2B heteromerization occurs primarily in the NAc and is maintained during cocaine withdrawal in the core subdivision.

### Cocaine-evoked potentiation of glutamate transmission onto D1R-MSNs requires D1R-GluN1 heteromerization

To study the function of the D1R-GluN1 heteromers in cocaine-induced adaptations, we designed an adeno-associated virus (AAV)-based strategy to disrupt heteromers in a spatially- and temporally-controlled manner (Fig. 2A). The AAV Tet-On-GluN1C1 allows a doxycycline (dox)-inducible bicistronic expression of the RFP reporter protein together with a peptide corresponding to the C1 cassette (D_864_-T_900_) of GluN1 that binds to D1R (*28*). This peptide blocks D1R-GluN1 interaction *in vitro*, while preserving the functions of individual D1R and NMDAR independently of their heteromerization (*31*). The control virus, Tet-On-GluN1C1∆, encodes a C1 cassette deleted of 9 amino acids that are required for electrostatic interactions between D1R and GluN1 (*38*). This mutated cassette does not interfere with D1R-GluN1 interaction *in vitro* (*31*). After stereotaxic injections in the NAc, the treatment with dox triggered a rapid and sustained expression of RFP (Fig. 2A). In naive mice, analysis of D1R-GluN1 proximity in RFP-positive neurons showed that Tet-On-GluN1C1 significantly reduced D1R-GluN1 PLA puncta when compared to the control AAV (Fig. 2B,C), indicating an efficient disruption of these heteromers *in vivo.* We also verified that blocking D1R-GluN1 heteromerization altered downstream cocaine-mediated signaling events (*20*, *31*), including GluN2B phosphorylation and extracellular-signal regulated kinase (ERK) pathway activation (Fig. S2), without compromising neuronal survival (Fig. 2D).

**Figure 2.**
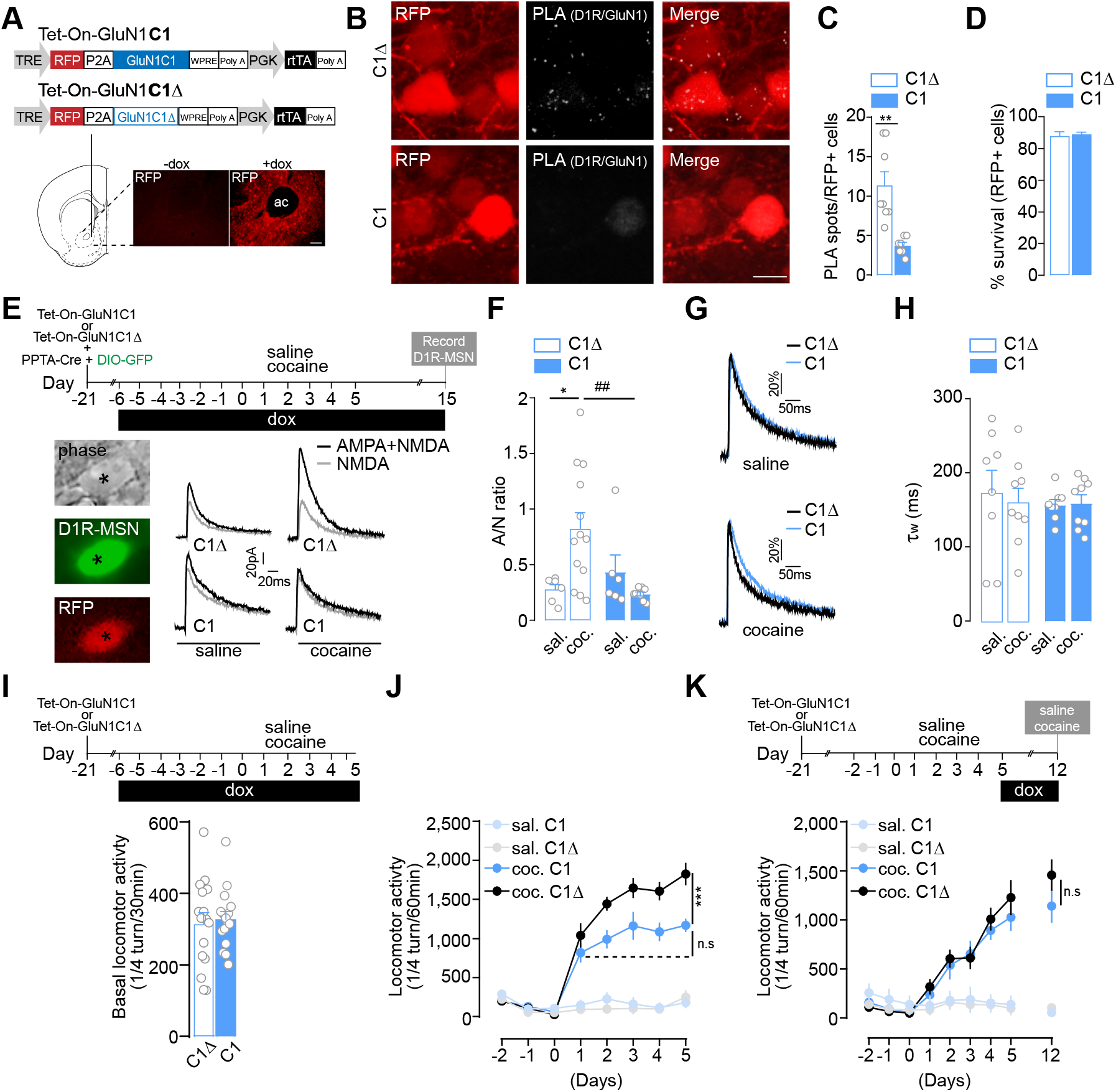
D1R-GluN1 heteromerization controls cocaine-evoked potentiation of glutamate transmission onto D1R-MSN and the development of behavioral sensitization. (**A**) Top: Viral-based strategy for expression of interfering peptide to disrupt D1R-GluN1 interaction (Tet-On-GluNC1; C1) and control (Tet-On-GluN1C1Δ; C1Δ) in the NAc. Bottom: Example image of doxycycline (+dox)-mediated RFP expression (ac: anterior commissure). (**B**) Example image of D1R-GluN1 heteromer detection by PLA in C1Δ- and C1-transduce neurons. Scale bar: 10 μm. (**C**) Quantifications of the PLA signal in C1Δ- and C1-transduced neurons. Two-sided Student’s t-test, *t* = 4.078 df. = 13, ** *P* =0.0013, n=7-8 cells from 4 mice/group. (**D**) Neuronal survival of C1Δ- and C1-transduced neurons. Two-sided Student’s t-test, *t* = 0.354 df. = 6, *P* =0.735, n=4 mice/group. (**E**) Experimental design and example trace of AMPA+NMDA (black) and NMDA (grey) currents in neurons (asterisk) expressing GFP(i.e. D1R-MSN (see Fig. S3A)), and C1 or C1Δ (RFP+). (**F**) AMPA to NMDA (A/N) ratios. Two-way ANOVA: virus effect, F(1, 30) = 2.511, *P = 0.033; ## P=0.0061; n= 3-4 mice/group and n=6-13 cells/group. (**G**) Comparison of representative recordings of pharmacologically-isolated NMDAR EPSCs, normalized to the peak amplitude (in %). (**H**) Deactivation kinetics of NMDAR EPSCs. Two-way ANOVA: virus effect, F (1, 30) = 0.205, P>0.999, n= 3-4 mice/group and n=8-9 cells/group. (**I**) Experimental time frame and basal locomotor activity. Two-sided Student’s t-test, *t* = 0.332 df. = 30, *P* =0.742, n=16 mice per group. (**J**) Inhibition of D1R-GluN1 heteromerization during the development of locomotor sensitization. Three-way ANOVA: virus effect,F(1, 256) = 13.72, ***P= 0.0003; n.s P>0.9999, n=7-11 mice/group. (**K**) Experimental time frame and measurement of locomotor activity in each group. Three-way ANOVA: virus effect, F (1, 243) = 0.6160, P>0.9999, n=7-8 mice/group. n.s not significant. Error bars denote s.e.m.

Long-lasting changes of glutamate transmission at cortical projections onto D1R-MSN of the NAc have been causally implicated in the development of cocaine-induced locomotor sensitization (*13*). To study the contribution of D1R-GluN1 heteromerization in drug-induced plasticity at these synapses, we injected mice with Tet-On-GluN1C1 together with a mixture of AAV-PPTA-Cre – driving the expression of the Cre recombinase under the control of the D1R-MSN-specific prepro-tachykinin promoter - and AAV-DIO-eGFP to tag D1R-MSN (*7*, *39*, *40*) (Fig. S3A). These mice were supplemented with dox before and during daily injections of saline or cocaine for 5 days (5 d) followed by 10 d of withdrawal. As previously shown (*16*), cocaine triggered an increase of AMPA/NMDA (A/N) ratio – an index of synaptic plasticity - in D1R-MSN of mice injected with the control virus in the NAc. By contrast, while preserving basal synaptic transmission in thesaline-treated group, the inhibition of D1R-GluN1 heteromerization blunted the cocaine-evoked increase in A/N ratio (Fig. 2E,F), without modifying the amplitude or the kinetics of NMDAR EPSCs (Fig. 2G,H). These data show that D1R-GluN1 heteromerization controls cocaine-evoked changes in glutamate transmission in D1R-MSNs.

### D1R-GluN1 heteromerization controls the development of cocaine-induced locomotor sensitization

We next evaluated the role of D1R-GluN1 heteromerization in the behavioral sensitizing properties of cocaine by supplementing Tet-On-AAV-injected mice with dox prior to and during saline or cocaine administration. Uncoupling D1R from GluN1 did not affect basal locomotion (Fig. 2I), nor the acute hyperlocomotor response triggered by the first cocaine injection, but fully blocked the development of the behavioral sensitization induced by subsequent injections (Fig. 2J). Of note, dox supplementation did not alter body weight, basal locomotion or behavioral sensitization to cocaine in mice that were not injected with Tet-On viruses (Fig. S4A-C).

By temporally controlling expression of the interfering peptides, we assessed the contribution of D1R-GluN1 heteromerization to the maintenance phase of locomotor sensitization. Mice injected with Tet-On viruses were treated with saline or cocaine for 5 d in the absence of dox. As expected, these mice displayed a similar cocaine-induced locomotor sensitization regardless of the virus injected. To switch off D1R-GluN1 heteromerization after behavioral sensitization, dox was given after the last saline or cocaine injection and during a 7 d withdrawal followed by a challenge injection of saline or cocaine (Fig. 2K-top).

We found that mice displayed the same level of sensitization in response to the cocaine challenge regardless of the AAV used, demonstrating that D1R-GluN1 heteromerization is not required for the maintenance of locomotor sensitization (Fig. 2K-bottom). Since cocaine also enhanced D1R-GluN1 heteromerization in the dorsal striatum (Fig. 1C), we also targeted heteromers in this striatal sub-region and obtained the same results (Fig. S4D,E). Overall, these data show that D1R-GluN1 heteromerization in the striatum controls the development, but not the maintenance, of cocaine’s sensitizing effects.

### D2R-GluN2B heteromerization selectively controls the maintenance of cocaine sensitizing effects

In light of the significant and persistent impact of cocaine on D2R-GluN2B heteromerization in the NAc (see Fig. 1D,E), we generated AAV Tet-On-D2R-IL3 to achieve a dox-inducible expression of a peptide corresponding to a small fragment (T_225_-A_234_) located within the 3rd intracellular loop (IL3) of D2R. This IL3 domain is known to play a key role for D2R-GluN2B interaction (*32*). Since critical amino acids responsible for D2R-GluN2B interaction have not been yet identified within this D2R-IL3 fragment, we used a control virus (Tet-On-D2R-IL3-scr) driving the expression of a scrambled peptide (Fig. 3A). The Tet-On-D2R-IL3 efficiently reduced D2R-GluN2B PLA puncta and preserved neuronal survival (Fig. 3B-D). Importantly, this peptide altered the interaction of D2R with GluN2B while sparing the functions of individual component receptors, as shown by its lack of effect on D2R-mediated inhibition of cAMP production (Fig. 3E) and NMDA currents (see below).

**Figure 3.**
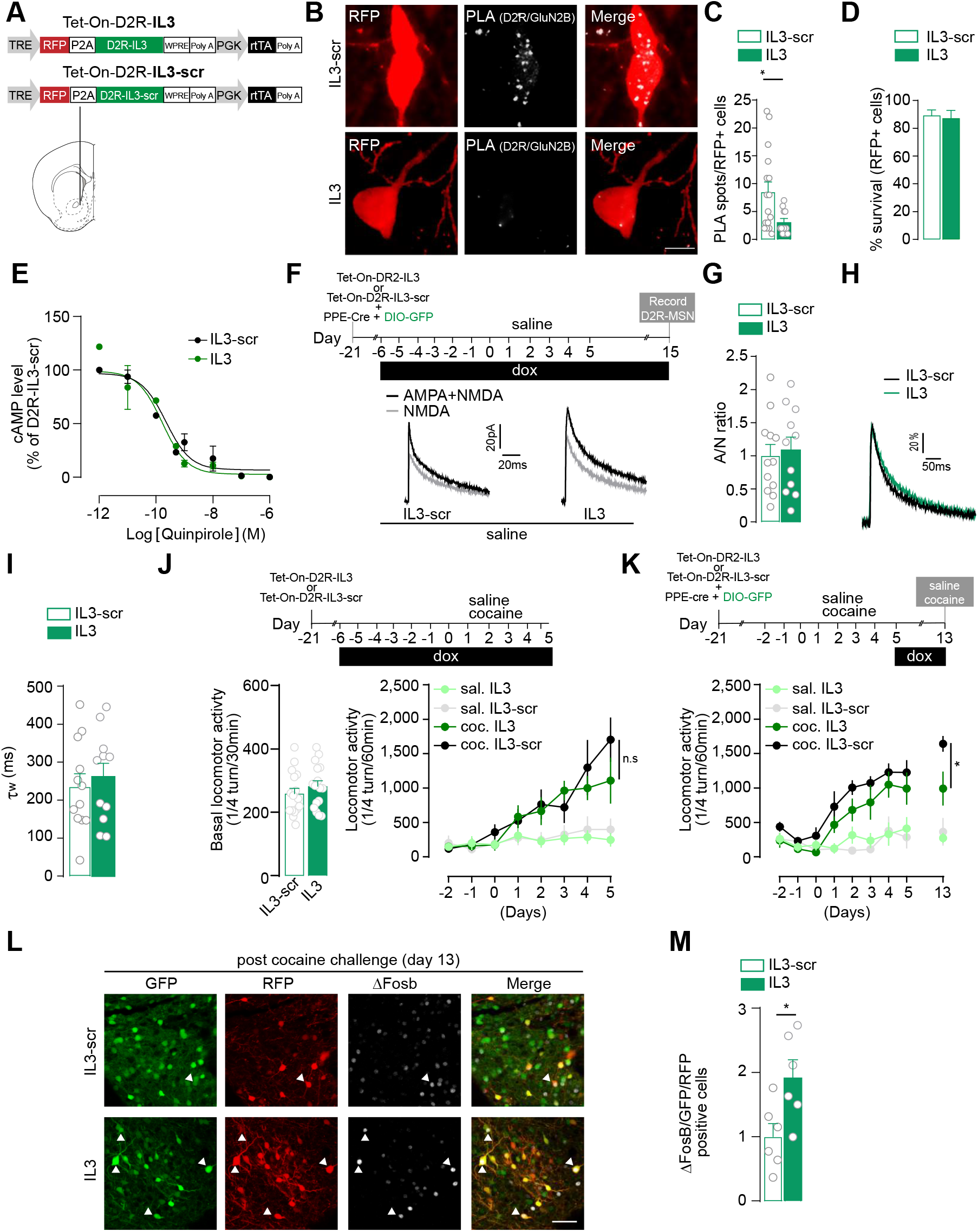
D2R-GluN2Bheter omerization selectively controls the maintenance of cocaine sensitizing effects. (**A**) Viral strategy for expression of interfering peptide to prevent D2R-GluN2B heteromerization(Tet-On-D2R-IL3; IL3) and control (Tet-On-D2R-IL3-scr; IL3-scr). (**B**) Representative image D2R-GluN2B heteromer detection by PLA in IL3-scr- and IL3-transduced neurons. Scale bar: 10 μm. (**C**) Quantifications of the PLA signal in IL3-scr- and IL-3 infected neurons. Two-sided Student’s t-test, t = 2.393 df. = 25, **P =0.0246, n=11-16 cells from 4 mice/group. (**D**) Neuronal survival of IL3-scr- and IL3-transduced neurons. Two-sided Student’s t-test, t = 0.2767 df. = 6, P =0.7913, n=4 mice/group. (**E**) Tet-On-D2R-IL3 spares quinpirole-induced inhibition of forskolin-induced accumulation of cAMP. LogIC_50_ is −9.78 for D2R-IL3 and −9.65 for D2R-IL3-Scr. n=3 independent experiments/condition. (**F**) Experimental time frame and representative traces of AMPA+NMDA (black) and NMDA (grey) currents in neurons expressing GFP (i.e. D2R-MSN (see Figure S3B)) and IL3 or IL3-scr (RFP+). (**G**) AMPA to NMDA (A/N) ratios; Two-sided Student’s t-test, t = 0.3397 df. = 21, P =0.7375, n=4-6 mice/group and n=11-12 cells/group. (**H**) Comparison of representative recordings of pharmacologically-isolated NMDAR EPSCs, normalized to the peak amplitude (in %). (**I**) Deactivation kinetics of NMDA EPSCs, Two-sided Student’s t-test, t = 0.5129 df. = 21, P =0.6134, n=11-12 cells/group. (**J**) Top: Experimental time frame. Bottom left: basal locomotor activity; Two-sided Student’s t-test, t = 0.994 df. = 30, P =0.3282, n=16 mice/group. Bottom right: Inhibition of D2R-GluN2B heteromerization does not impair the development of locomotor sensitization Three-way ANOVA: virus effect, F (1, 192) = 1.984, n.s P >0.9999; n=6-8 mice/group. (**K**) Experimental time frame and impact of D2R-GluN2B heteromer inhibition on the maintenance of cocaine-evoked locomotor sensitization. Three-way ANOVA: virus effect, F (1, 198) = 7.278, * P = 0.0330; n=6 mice/group. (**L**) Representative images of ΔFosB expression in D2R-MSN (GFP+) infected with IL3-scr or IL3 (RFP+) after the cocaine challenge injection (see panel K). Arrowheads show GFP+/RFP+/ ΔFosB+ D2R-MSN. Scale bar: 50 μm. (**M**) Quantifications of GFP+/RFP+/ ΔFosB+ D2R-MSN. Two-sided Student’s t-test, t = 2.694 df. = 10, *P =0.0225, n=6 mice/group. n.s: not significant. Error bars denote s.e.m.

Since repeated cocaine exposure does not modify A/N ratio in D2R-MSN (*16*), the consequences of uncoupling D2R from GluN2B was studied in saline-treated animals with virally-tagged D2R-MSN owing to the co-injection of AAV-PPE-Cre – driving the expression of the Cre recombinase under the control of the D2R-MSN-specific prepro-enkephalin promoter (*7*) and AAV-DIO-eGFP (Fig. S3B). We found that the inhibition of D2R-GluN2B heteromerization did not alter, by itself, A/N ratio in D2R-MSN when compared to D2R-MSN transduced with the control virus (Fig. 3F,G). Of note, Tet-On-D2R-IL3 also left unchanged the amplitude and kinetics of NMDA currents (Fig. 3H,I), indicating a lack of non-specific effect on individual NMDAR functions, thereby validating our interfering viral strategy.

At the behavioral level, interfering with D2R-GluN2B heteromerization preserved both basal locomotion (Fig. 3J-left) and the development of cocaine-induced locomotor sensitization (Fig. 3J-right).

Strikingly, we found that the inhibition of D2R-GluN2B heteromerization during cocaine withdrawal reduced the maintenance of the behavioral sensitization compared to cocaine-treated mice injected with the control virus (Fig. 3K). These mice were sacrificed to analyze ΔFosB expression levels, used here as a proxy for neuronal activity. We observed a significant increase of D2R-MSN expressing ΔFosB after the cocaine challenge upon inhibition of D2R-GluN2B interaction (Fig. 3L,M). Since D2R-Glu2NB heteromerization has been shown to mediate the D2R agonist-induced inhibition of NMDAR *ex vivo* (*32*), our data suggest that the impaired maintenance of the sensitizing effects of cocaine observed upon D2R-GluN2B uncoupling may result from increased D2R-MSN activity.

Altogether, these data demonstrate that D1R-GluN1 and D2R-GluN2B heteromerization controls the development and maintenance of cocaine sensitizing effects, respectively.

### Differential roles of D1R-GluN1 and D2R-GluN2B heteromers in the rewarding effects of cocaine

We next investigated the role of these heteromers in the rewarding effects of cocaine using a conditioned place preference (CPP) paradigm. Mice injected in the NAc with a control virus and supplemented with dox developed a significant cocaine-induced CPP, which was blunted when D1R and GluN1 were uncoupled (Fig. 4A). Although single or repeated cocaine administration has been shown to trigger dendritic spine formation in D1R-MSN (*39*, *41*), there is no study showing such morphological changes in the NAc in the context of CPP. Mice were thus sacrificed the day after the behavioral test to perform a 3D morphological analysis of GFP-tagged D1R-MSN. Control mice, which developed CPP to cocaine, displayed a significant increase of dendritic spine density in D1R-MSN, which was inhibited when D1R and GluN1 were uncoupled (Fig. 4B). To study the implication of D1R-GluN1 heteromerization on relapse to CPP, AAV-injected mice were initially trained for CPP in the absence of dox. Once mice developed CPP, dox was added to alter D1R-GluN1 interaction during an extinction period followed by a cocaine-induced relapse. We observed that the inhibition of D1R-GluN1 interaction did not alter the kinetics of extinction, nor the relapse to CPP (Fig. 4C), thus supporting a critical role for D1R-GluN1 heteromers in the development of the rewarding effects of cocaine, as observed for locomotor sensitization, but not in the propensity to relapse.

**Figure 4.**
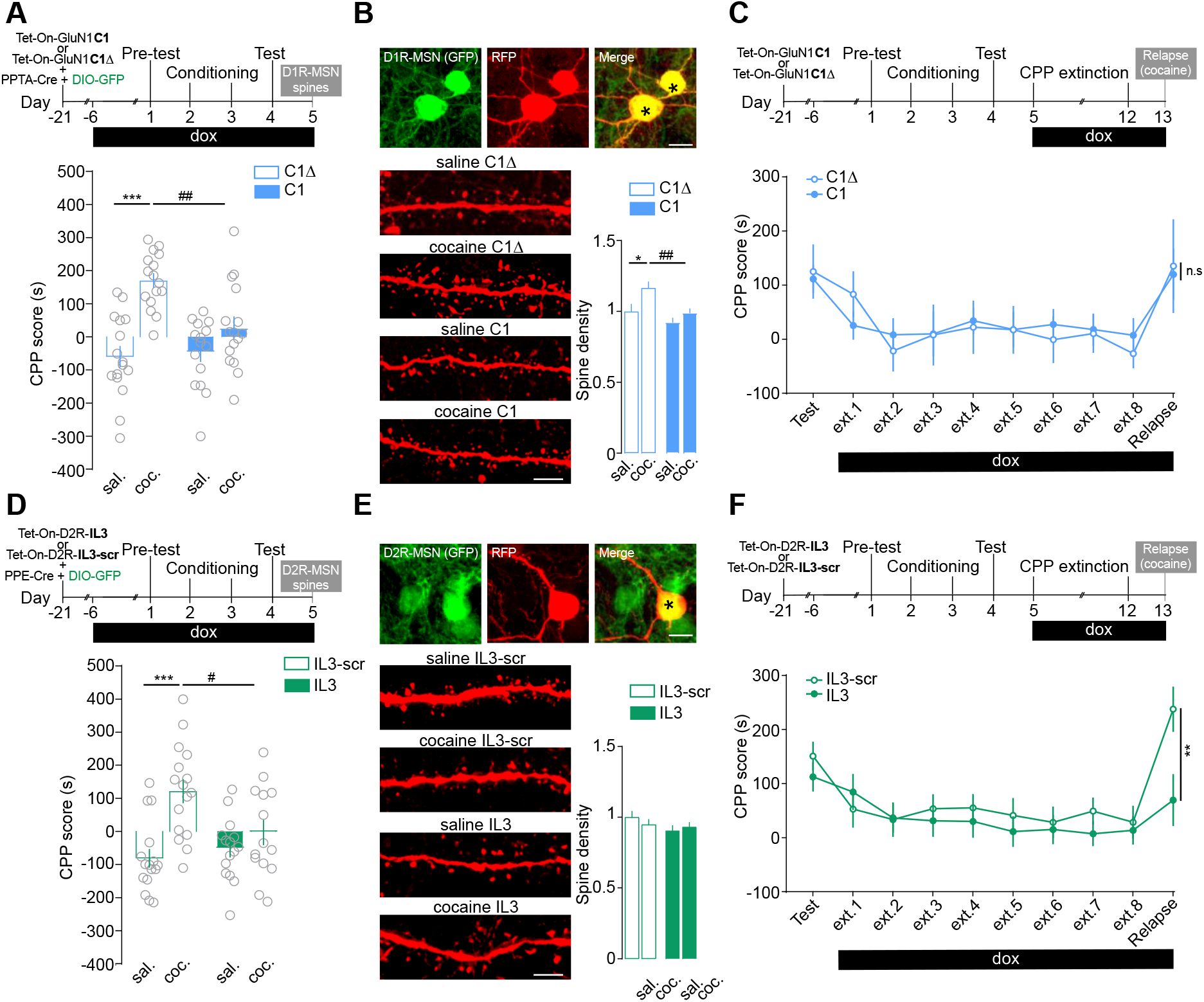
Differential roles of D1R-GluN1 and D2R-GluN2B heteromerization in controlling the rewarding effects of cocaine. (**A**) Experimental time frame and conditioned place preference (CPP) score upon inhibition of D1R-GluN1 heteromerization. Two-way ANOVA: virus effect, F(1, 59) = 5.281,*** P<0.0001; ^##^P=0.0040, n=15-16 mice/group. (**B**) Top: low magnification images of D1R-MSN (GFP+; see Fig. S3A) infected (RFP) shown by the asterisks Scale bar 10 μm. Bottom: high magnification of dendritic segments. Scale bar: 5 μm. Spine density analysis. Two-way ANOVA: virus effect, F(1,162) = 11.14, * P=0.0446;## P=0.0043,n=27 −69 dendrites from 6 mice/group. (**C**) Experimental time frame to study the impact of D1R-GluN1 uncoupling on the extinction and cocaine induced-relapse to CPP. Two-way ANOVA: virus effect, F (1, 24) = 0.004, P > 0.999, cocaine C1 vs cocaine C1Δ, CPP score on relapse day, n=10-16 mice/group. (**D**) Same as for (A) upon inhibition of D2R-GluN2B heteromerization. Two-way ANOVA: virus effect, F (1, 57) = 2.424, ^***^P <0.0001; ^#^P=0.0396, n=14-16 mice per group. (**E**) Top: low magnification images of D2R-MSN (GFP; see Fig S3B) infected (RFP) shown by the asterisk. Scale bar 10 μm. Bottom: high magnification of dendritic segments. Scale bar: 5 μm. Spine density analysis. Two-way ANOVA: virus effect, F (1, 166) = 0.1268, n.s P >0.999 saline IL3-scr vs cocaine IL3-scr; n=30-53 dendrites from 6 mice/group. (**F**) CPP score upon inhibition of D2R-GluN2B during CCP during extinction and cocaine-induced relapse. Two-way ANOVA: virus effect, F (1, 31) = 0.899, ^**^P =0.0018, cocaine IL3 vs cocaine IL3-scr CPP score on relapse day, n=16-17 mice/group. n.s: not significant. Error bars denote s.e.m.

The uncoupling of D2R from GluN2B also blocked the development of cocaine CPP (Fig. 4D) but this was not correlated to morphological changes in D2R-MSNs since the CPP paradigm did not trigger any modification of dendriticspine density in D2R-MSN regardless of the AAV used (Fig. 4E). Inhibiting D2R-GluN2B heteromerization once the mice have developed CPP did not impact the extinction of CPP but significantly reduced cocaine-induced relapse (Fig. 4F). This indicates that D2R-GluN2B heteromerization is required for both the development and relapse of cocaine-induced CPP, independently of morphological changes in GFP-tagged D2R-MSNs.

### Inhibiting D1R-GluN1 or D2R-GluN2B heteromerization does not alter conditioned place preference for food

Manipulating D1R-GluN1 or D2R-GluN2B heteromerization *in vivo* allowed us to reveal their selective implication in controlling distinct phases of long-term cocaine-evoked adaptations. We next examined whether receptor heteromerization also controls non-drug reward processing. We found that disrupting either heteromer subtype, using comparable conditions as for our studies with cocaine, failed to alter the rewarding properties of food (Fig. 5), supporting a role of these heteromers in controlling the rewarding effects of cocaine but not a non-drug reward.

**Figure 5.**
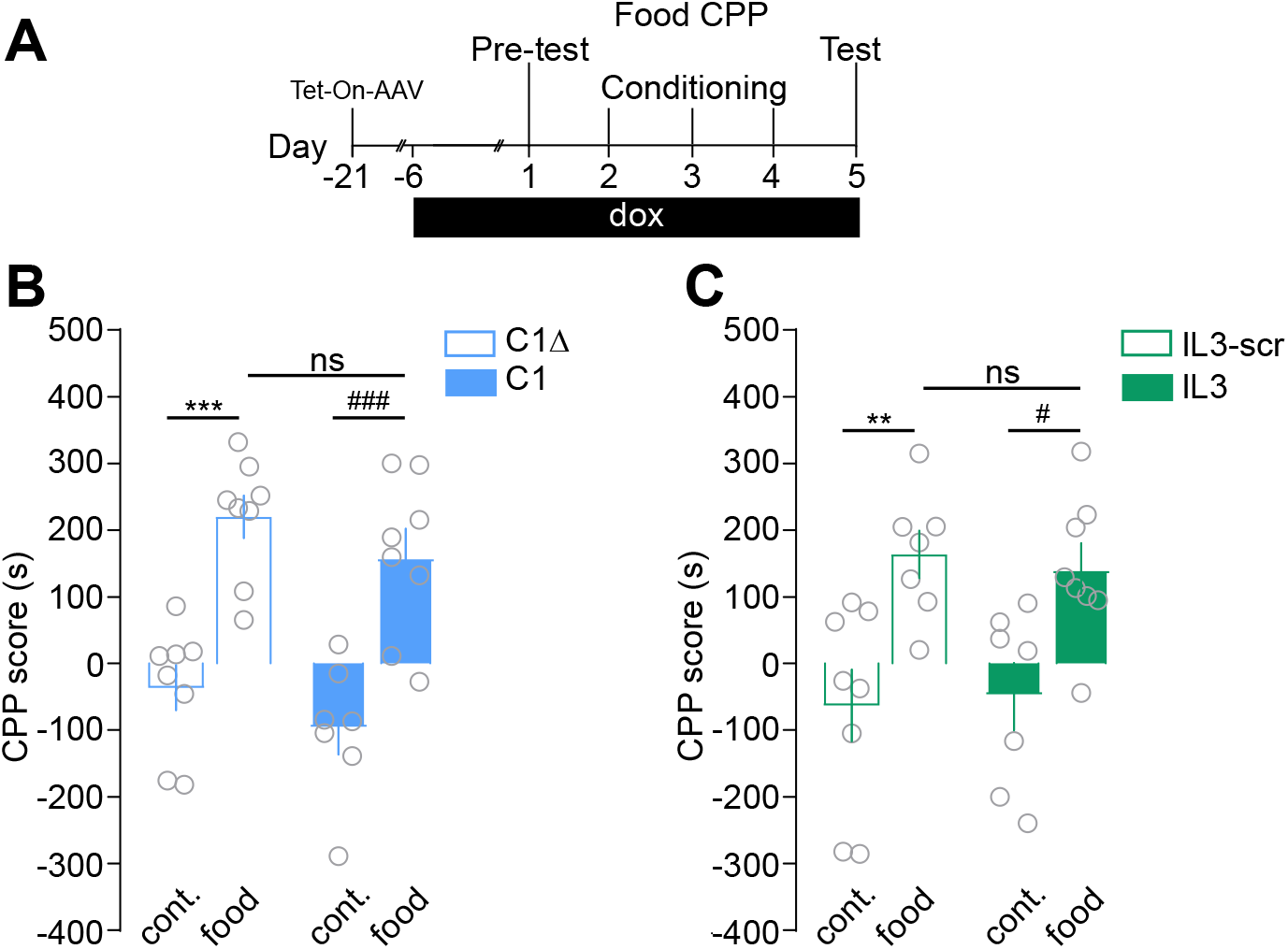
Inhibiting D1R-GluN1 or D2R-GluN2B heteromerizationdoes not alterconditionedplacepreference for food. (**A**) Experimental time frame to study the consequences of inhibition of heteromerization on the development of food-induced CPP. (**B**) Impact of inhibiting D1R-GluN1 heteromerization on the CPP score. Two-way ANOVA: virus effect, F (1, 27) = 2.756, ***P =0.0002, control C1Δ vs food C1Δ; ^###^P =0.0002, control C1 vs food C1; n.s P>0.999, food C1 vs food C1Δ, n=7-8 mice/group. (**C**) Effect of inhibiting D2R-GluN2B heteromerization on the CPP score. Two-way ANOVA: virus effect F (1, 26) = 0.007, **P <0.0098, control IL3-scr vs food IL3-scr; #P =0.0366, control IL3 vs food IL3; n.s, P>0.999 food IL3 vs food IL3 -scr, n=7-8 mice/group. n.s: not significant. Error bars denote s.e.m.

### D2R-GluN2B heteromerization is increased in post-mortem brain samples from addict subjects despite decreased D2R expression

As evidenced above, D2R-GluN2B heteromerization plays a cardinal role in the maintenance of cocaine’s effects without affecting natural reward processing in mice, which positions this heteromer subtype as a potential therapeutic target for drug addiction. We therefore investigated whether D2R-GluN2B heteromerization could be detected in human brain tissues and modulated in subjects with a history of psychostimulant dependence.

PLA has recently been shown as a suitable approach to detect single proteins or receptor heteromers, including D2R-A2AR, in human brain samples (*42*, *43*). We therefore performed single detection of D2R, GluN2B and D2R-GluN2B heteromers in post-mortem human samples from control subjects and matched individuals with a history of dependence. Although often poly-addicts, these individuals were selected for their main dependence to psychostimulants and the presence of traces of psychostimulants in their blood at the time of death (Table S1). From whole-slide images of caudate putamen samples, automated detection of PLA signal was performed from 25 high-magnification images per subjects randomly selected within the ventral part of the samples, which corresponds to the mouse NAc (Fig. S5).

D2R single detection produced a dense punctate pattern in control subjects (Fig. 6A), as already reported (*42*). As expected, this signal was absent when PLA was performed in the absence of the primary antibody. Interestingly, we detected a significant decrease of relative D2R protein expression in sample from addicts compared to control subjects (Fig. 6B), consistent with the well-established decrease of striatal D2R availability reported in psychostimulant abusers by PET imaging (*44*–*48*). By contrast, GluN2B single detection produced a dense GluN2B signal that was not different between samples from addicts and controls (Fig. 6C,D). The double recognition of D2R-GluN2B proximity yielded a punctate signal in control subjects, which was undetectable when one of the two primary antibodies was omitted (Fig. 6E).

**Figure 6.**
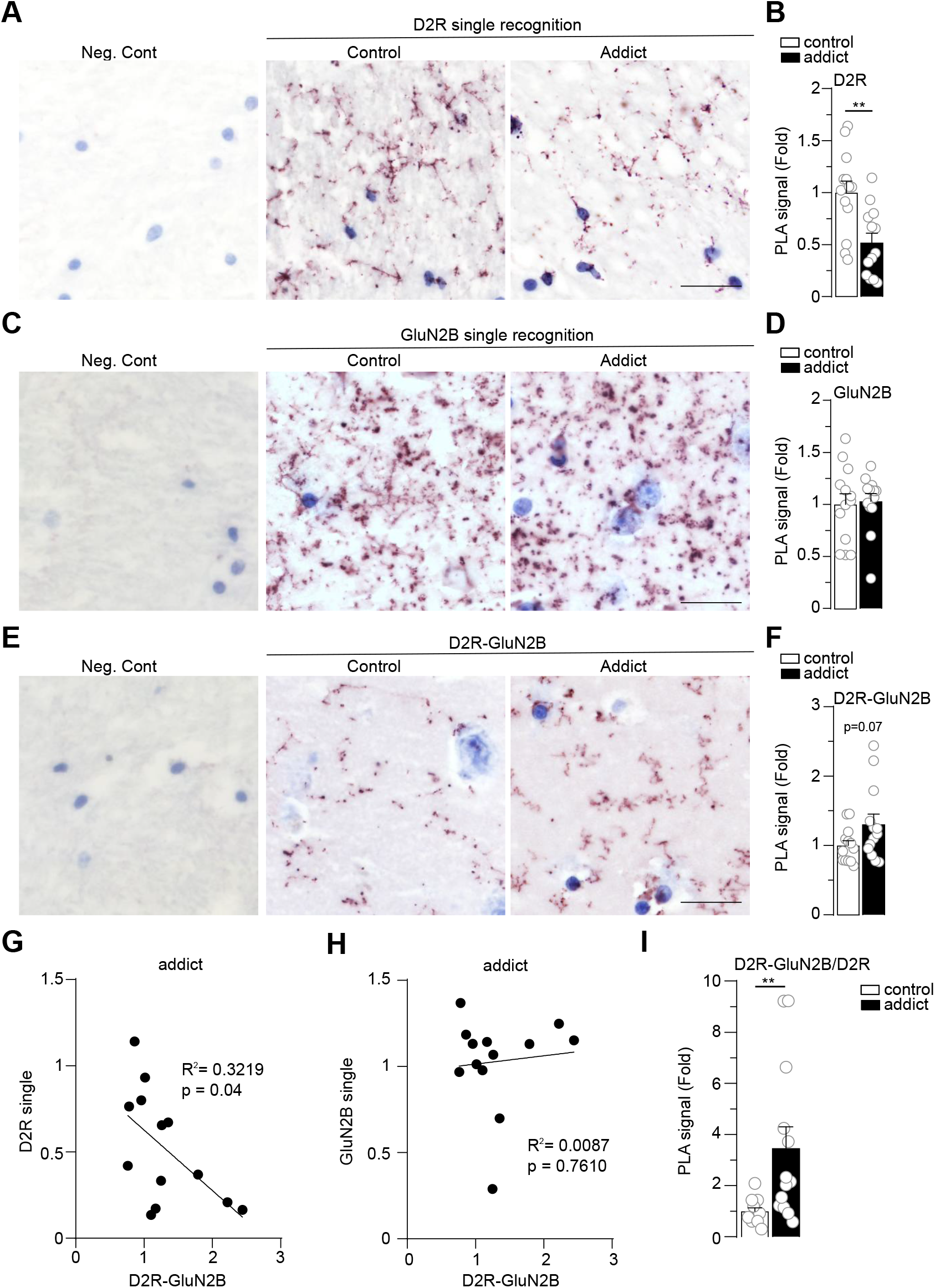
D2R-GluN2B heteromerization is increased in post-mortem brain samples from addict subjects despite decreased D2R expression. (**A**) Representative images of D2R single recognition by PLA and negative control, in which the primary antibody is omitted (left panel Neg. Cont; see Fig. S5). (**B)** Quantifications of D2R single PLA signal represented as fold decreased compared to control subjects. Two-sided Student’s t-test, t = 3.331 df. = 24, **P =0.0028, n=13 subjects/group. (**C**) Example images of GluN2B single detection and Neg. Cont. (**D**) Quantifications of Glu2NB PLA signal. Two-sided Student’s t-test, t = 0.224 df. = 24, P =0.8243, n=13 subjects/group. (**E**) Illustrative images of D2R-GluN2B heteromer detection by PLA and Neg. Cont (GluN2B antibody omitted). (**F**) Quantifications of D2R-GluN2B PLA signal. Two-sided Student’s t-test, t = 1.868 df. = 24, P =0.074, n=13 subjects per group. (**G**) Pearson correlation between D2R expression levels and D2R-GluN2B heteromerization for each sample from all addict subjects, R²=0.3219, P=0.0432. (**H)** Pearson correlation between GluN2B expression levels and D2R-GluN2B heteromerization for each sample from all addict subjects R^2^=0.0087, P=0.761. (**I**) Quantifications of D2R-GluN2B PLA signal normalized to D2R expression levels for each subject. Two-sided Student’s t-test, t = 2.882 df. = 24, **P =0.0082, n=13 subjects/group. (A, C, E) Scale bar: 25 μm. Error bars denote s.e.m.

In samples from addicts, there was a trend towardsan increase of D2R-GluN2B heteromers in samples from addicts (Fig. 6F), despite the drastic decrease of D2R levels. We therefore analyzed whether a correlation between D2R levels and D2R-GluN2B heteromerization could exist within each addict sample. We found a significant inverse correlation between these two parameters (Fig. 6G), suggesting that despite the lower levels of D2R expression in addicts, at least a remaining pool of D2R was preferentially involved in D2R-GluN2B heteromerization. By contrast, there was no correlation between GluN2B expression and D2R-GluN2B heteromerization (Fig. 6F). Considering the decreased D2R levels in samples from addicts, we normalized the D2R-GluN2B PLA signal to D2R levels for each individual and found a significant increase in relative D2R heteromerization with GluN2B in samples from addicts when compared to controls (Fig. 6I). To the best of our knowledge these results provide the first evidence of a decreased D2R protein expression in thestriatum of psychostimulant addicts, which is associated with an increase of D2R-GluN2B heteromerization. Together with our interventional approach in mice, our data support D2R-GluN2B heteromers as therapeutic targets of potential interest in addiction.

## Discussion

Optogenetic studies undeniably showed that distinct phases of drug-induced behavioral adaptations rely on DA-evoked synaptic adaptations at specific glutamate inputs onto MSN subpopulations (*13*-*15*, *49*, *50*). Nonetheless, the underlying molecular mechanisms remain poorly understood (*2*). This is an important issue because the identification of events responsible for such a detrimental interplay between dopamine and glutamate signaling may help in the development of innovative strategies with therapeutic potential. Herein, we provide multiple lines of evidence, from mice to humans, that the heteromerization of glutamate NMDAR with D1R or D2R is enhanced by psychostimulants and preferentially controls the development and maintenance phases of cocaine-evoked long-term adaptations, respectively.

The focus on dopamine and NMDA receptor heteromers as potential integrators of dopamine and glutamate inputs that may control drug-mediated adaptations stems from *in vitro* and *ex vivo* studies showing that such a direct physical interaction allows a reciprocal fine-tuning of the component receptors’ functions (*23*, *27*). In particular, patch-clamp recording from striatal slices showed that D1R-GluN1 and D2R-GluN2B interactions respectively facilitate and inhibit NMDAR-mediated signaling upon DA increase (*31*, *32*). An appealing hypothesis would therefore be that, by linking dopamine to glutamate signaling in opposite ways, these heteromers could constitute molecular substrates for drugs of abuse to exert their differential effects on the activity of MSN subtypes, which has been proposed to underlie the switch from recreational drug consumption to addiction (*3*, *51*).

In agreement with this model, our PLA analysis showed that locomotor sensitization induced by repeated cocaine injections was associated with an increase of both heteromers in the NAc. We also found that this increased heteromerization requires dopamine receptor stimulation. While the PLA method cannot establish the direct physical contact of the two proteins or the stoichiometry of the complex, it does indicate that the proteins are in close molecular proximity (*34*). Nonetheless, previous studies have provided evidence for direct interactions between the D1R-GluN1 and D2R-GluN2B (*28*, *32*) and we find that our viral minigenes selectively decrease the PLA signals; we therefore interpret these PLA data as support for receptor heteromerization. While in-depth characterization of the molecular events responsible for this increased receptor interaction upon cocaine exposure is beyond the scope of this study, a possible explanation may lie in the observation that repeated cocaine exposure decreases PSD-95 expression in the NAc (*36*). In fact, PSD-95 is a known endogenous inhibitor of D1R-GluN1 interaction (*37*) that also binds to D2R (*52*) and the GluN2B c-terminal end (*53*). Even though the PLA approach is able to provide a snapshot of the impact of cocaine on receptor heteromerization *in situ* in their native environment (*31*, *34*, *35*), future work is needed to investigate whether heteromerization of DA and NMDA receptors is an input-specific process and whether it relies on the modulation of receptor surface expression and/or dynamics.

The development of the sensitizing effects of cocaine has been previously causally linked to the potentiation of glutamate transmission at cortical projections onto D1R-MSN of the NAc (*13*). Since we observed that preventing D1R-GluN1 heteromerization reversed both alterations in A/N ratio and the development of behavioral sensitization, while sparing the function of individual component receptors (see (*31*)), our results suggest that D1R-GluN1 heteromers are key molecular platforms for the development of cocaine-induced long-term adaptations. In contrast, disrupting D1R-GluN1 interaction during a withdrawal from cocaine did not impact maintenance of the sensitized state, supporting a preferential role of this heteromer subtype in the initial phases of cocaine-mediated adaptations. In agreement with this hypothesis, we observed that preventing D1R-GluN1 heteromerization during CPP conditioning also blocked the development of the rewarding effects of cocaine, but failed to alter the extinction and relapse phases. This critical time window of D1R-GluN1 heteromer function restricted to the early developmental phase of cocaine-evoked adaptations agrees with our observation that cocaine-induced D1R-GluN1 heteromerization is a transient mechanism that does not outlast a 7 d withdrawal period. Instead, the temporally-controlled disruption of D2R-GluN2B heteromers revealed their preferential role in the maintenance of the sensitizing and rewarding effects of cocaine. In agreement with these findings, we found that D2R-GluN2B heteromerization persisted through withdrawal from cocaine. Moreover, this persistent heteromerization was specifically observed in the NAc core, which has been identified as a common output structure of neuronal circuits involved in both cue- and drug-induced relapse (*54*). Since the optogenetic activation of D2R-MSNs in the NAc has been shown to preserve cocaine-induced locomotor sensitization but to blunt its expression after withdrawal (*55*), our results support a model by which inhibiting endogenous D2R-GluN2B interaction during withdrawal hinders the persistence of cocaine-evoked responses by potentiating D2R-MSN activity. In support of this hypothesis, we found that the alteration of the maintenance phase of locomotor sensitization observed upon D2R-GluN2B heteromer disruption was associated with an increase of D2R-MSN activity, as revealed by an increased expression of Δ FosB in D2R-MSN. These observations therefore suggest that D2R-GluN2B heteromerization is a key molecular mechanism triggered by cocaine that dampens D2R-MSN activity and contributes to the persistence of cocaine-evoked adaptations.

Direct manipulations of MSN activity have clearly revealed that D1R-MSN and D2R-MSN activation, respectively, facilitates and blunts the development and maintenance phases of psychostimulant-induced behavioral adaptations (*2*). Strikingly, our findings that D1R-GluN1 and D2R-GluN2B heteromerization are involved in the induction and maintenance of cocaine-induced locomotor sensitization, respectively, highlight that these receptor complexes mediate discrete properties of MSN subpopulations and play complementary roles to mediate the full panel of cocaine-induced adaptations. Importantly, in further support of specific functions of dopamine and glutamate receptor heteromers, we established that their roles in shaping reward processing depends on the nature of the reward, since the disruption of either receptor heteromer blocked the development of cocaine-induced CPP but spared food-mediated CPP. Although the mechanisms underlying such selectivity to drug reward remain to be established, our findings suggest that targeting dopamine-glutamate receptor heteromers has the potential to preferentially alleviate pathological adaptations induced by drugs of abuse. In particular, the role for D2R-Glu2NB heteromerization in maintaining cocaine-induced adaptations combined with its lack of implication in reward processing to a natural reinforcer suggest that D2R-GluN2B heteromers are targets of choice from a translational standpoint. This led us to investigate whether this heteromer subtype could be detected in post-mortem human samples and modulated in subjects with a history of psychostimulant addiction.

Our PLA analysis revealed a strong reduction of D2R protein levels in the NAc of drug abusers. With the “single PLA” approach, the polyclonal secondary antibodies can bind to either a single primary antibody – therefore detecting a single antigen on the D2R – or to two different primary antibodies bound to proximal antigens – potentially revealing D2R homodimers. However, the latter is likely to be much less efficient and we assume that the single PLA signal in our study mainly reflects the density of single D2R (*42*). This first observation of a decreased D2R protein expression in post-mortem brain samples from addicts is consistent with the downregulation of D2R mRNA levels that has been described after long-term cocaine exposure in rats (*56*). Importantly, this finding could also partly account for the decrease in D2R binding readily observed with PET imaging of the striatum of drug abusers (*44*–*48*). Despite such downregulation of D2R protein, the proportion of D2R forming heteromers with GluN2B was three-fold higher in psychostimulant abusers compared to healthy subjects. Strikingly, addict individuals bearing the lowest D2R expression displayed the highest density of D2R-GluN2B. This raises questions regarding the underlying molecular mechanism of D2R-GluN2B formation in response to psychostimulant exposure in human. Based on our findings in mice that cocaine-induced D2R-GluN2B heteromerization depends on D2R stimulation, it is tempting to speculate that repeated increases of phasic dopamine levels resulting from recurrent psychostimulant consumption by addict individuals could be responsible for the higher D2R-GluN2B receptor proximity. The increased formation of D2R-GluN2B heteromerization we observed in human samples from psychostimulant abusers, together with interventional approaches in mice, emphasize their roles in the persistence of cocaine’s behavioral effects. These important findings constitute a significant breakthrough in understanding of the molecular bases of cocaine-induced adaptations and highlight the potential benefit of targeting D2R-Glu2NB heteromerization, not only in the field of addiction, but also potentially for multiple neuropsychiatric disorders associated with an imbalance of DA and glutamate transmission.

## Materials and methods

### Animals

6-week-old C57BL/6J male mice were purchased from Janvier labs (Le Genest, St Isle, France). The animals were housed four per cage, in a 12-hour light-dark cycle, in stable temperature (22°C) and humidity (60%) conditions with *ad libitum* access to food and water. They were acclimatized to the animal facility for at least 1 week. All experiments were carried out in accordance with the standard ethical guidelines (European Community Council Directive on the Care and Use of Laboratory Animals (86/609/EEC) and the French National Committee (2010/63)).

### Drugs

Drugs were administrated intraperitoneally in a volume of 10 ml/kg. Cocaine hydrochloride (Sigma Aldrich, St. Louis, MO) was dissolved in a saline solution (0.9% NaCl w/v).

9-tert-butyl doxycycline hydrochloride (9-TB-dox; Tebu-bio, Le Perray-en-Yvelines, France) was dissolved in a saline solution containing DMSO (5%) and Tween20 (5%). SCH23390 (0.25 mg/kg) or Eticlopride (0.5 mg/kg) dissolved in a saline solution (0.9% NaCl w/v) were administered 30 min prior to the challenge cocaine injection.

### Viral constructions

All AAV recombinant genomes were packaged in serotype 9 capsids. AAV-Tet-On-GluN1C1 expresses bicistronically the fluorescent reporter protein RFP and the C1 cassette of the GluN1 subunit (_864_DRKSGRAEPDPKKKATFRAITSTLASDT_900_) upon doxycycline (dox) treatment. The related control virus AAV-Tet-On-GluN1C1Δ expresses a truncated version of C1 that is deleted from a stretch of 9 positively charged amino acids (_890_S890FKRRRSSK_898_), which are required for D1R-GluN1 interaction55. The AAV-Tet-On-D2R-IL3 encodes a sequence of the third intracellular loop of the D2R (_225_TKRSSRAFRA_234_) interacting with GluN2B. The control 9AAV-Tet-On-D2R-scr expresses a scrambled sequence (KFARRTSASR) of the D2R-IL3 (full AAV sequences are available upon request). All Tet-On AAV were injected bilaterally by infusing 0.7 μl of a solution at 5.10^13^ viral genomes/ml per hemisphere for the NAc (2 μl for the dorsal striatum). The AAV-PPTA-Cre and AAV-PPE-Cre contain an expression cassette consisting of the Cre recombinase driven by the promoter of the PPTA gene (prepro-tachykinin) or the PPE gene (preproenkephalin), which are specifically expressed in D1R-MSN and D2R-MSN, respectively (*7*, *39*, *40*) (see supplementary Fig. 3). AAV PPTA-cre or AAV-PPE-cre were co-injected with the AAV-pCAG-DIO-eGFP-WRPE (Upenn) expressing flexed eGFP under the CMV/actin hybrid promoter (CAG). All viruses were diluted in PBS pluronic 0.001%.

### Stereotaxic injections

Mice were anesthetized with ketamine (150 mg/kg) and xylazine (10 mg/kg) and placed on a stereotaxic apparatus (David Kopf Instruments, Tujunga, CA, USA). Craniotomies were realized using the following coordinates: 1.7 mm rostral to the bregma, 1.2 mm lateral to midline and 4.6 mm ventral to the skull surface to target the NAc and 1mm rostral to the bregma, 1.8 mm lateral to midline and 3.25 mm ventral to the skull surface for the dorsal striatum. Viral injections were performed bilaterally at a rate of 0.15 μl/min using a 10 μl-syringe (Hamilton 1700 series, Phymep, Paris, France) with a 200 μm gauge needle (Phymep, Paris, France) mounted on a microinfusion pump (Harvard Apparatus, Holliston, MA). After the injection, the needle was left in place for an additional 8 min to avoid backflow.

### Doxycycline treatments

Three weeks after stereotaxic injections of Tet-On AVV, the expression of the constructs was triggered by daily intraperitoneal (IP) injection of 9TB-dox (10 mg/kg) for 4 days (4d). To maintain expression mice were then supplemented with a mix containing doxycycline (dox) Hcl (2 mg/ml), 9TB-dox Hcl (80μg/ml) and sucrose (1%) added in drinking water.

### Behavioral testing

All behavioral tests were conducted during the light phase (8:00–19:00). Animals were randomly assigned to the saline or cocaine groups after viral injection. Prior to behavioral testing, mice were handled daily during 7d in the experiment room. All mice were perfused with 4% (w/v) paraformaldehyde (PFA) 24h post-behavior to systematically verify the accuracy of stereotaxic injections and expression of the RFP reporter protein. Mice that did not meet quality criterion (i.e. non-bilateral expression, off-target diffusion, excessive backflow or low RFP expression) were discarded from the study.

### Locomotor activity and cocaine psychomotor sensitization

Locomotor activity was measured in a low luminosity environment inside a circular corridor (Immetronic, Pessac, France) containing four infrared beams placed at each 90° angle. Locomotor activity was expressed as a cumulative count of crossings between quarters of the corridor for the indicated time. Mice were treated with dox 7d before and until the end of the experiment. Mice were habituated to the test apparatus for 3d; basal locomotor activity was recorded on the third day of habituation. Cocaine sensitization experiments consisted of five daily 90 min sessions during which spontaneous activity was recorded for 30 min before saline or cocaine (15 mg/kg) injections and locomotor activity was then measured for 60 min post-injections. To study the consequences of uncoupling DAR from NMDAR on the maintenance of cocaine-induced locomotor sensitization, mice were treated for 5 consecutive days with saline or cocaine in the absence of dox. After the last injections, mice were supplemented with dox during a withdrawal period followed by a challenge injection of saline or cocaine.

### Cocaine Conditioned Place Preference (CPP)

To study the impact of DAR-NMDAR heteromerization on the development of CPP, mice were treated with dox for 7d before and until the end of the experiment. The CPP was performed in a two-compartment Plexiglas Y-maze apparatus (Imetronic). Each compartment contains different visual cues and floor textures for which mice did not show any preference on average before conditioning. All sessions lasted 20min. On day 1, mice were placed in the center of the apparatus and allowed to explore freely both compartments. Time spent in each compartment was automatically recorded. Mice spending more than 70% of the time in one compartment were excluded. On day 2, to avoid any initial preference bias, mice were randomly assigned to one or the other compartment for each group. Mice were injected with saline and placed immediately in the assigned closed compartment for 20 min. After 1h, mice were injected with saline or cocaine and placed in the other closed compartment. This was repeated on day 3. The test was performed on day 4, during which mice had a free access to both chambers. The CPP score was calculated as the difference between the time spent in the cocaine-paired chamber during day 4 minus the time spent in this compartment on day 1. CPP extinction and maintenance experiments were performed on the cocaine groups of mice injected with Tet-On-AVV that developed a preference for the cocaine-paired chamber in the absence of dox. Mice were then treated with dox until of the behavioral assessment. For the extinction phase, mice were injected with saline and put back in the apparatus with free access to both compartments for 20 min daily for 8 days. On the ninth day, mice were injected with cocaine and allowed to explore both compartments.

For palatable food-induced CPP, mice were food-deprived to 90% of initial *ad libitum* weight and treated with dox 7d prior and during behavioral assessment. Experiments were performed in the same apparatus and conditions as for cocaine-induced CPP with the following modifications: On day 2, after random group assignment. Mice were placed immediately in the assigned closed compartment containing chocolate crisps (Chocapic, Nestlé, Vevey, Switzerland) or nothing for 20 min. After 1 h, mice were placed in the other closed compartment. This was repeated on day 3 and 4. The test was performed on day 5.

### Mouse tissue preparation

Mice were anesthetized with an I.P injection of Euthasol (100 mg/kg; Le Vet, Oudewater, Netherlands) and perfused transcardially with 0.1 M Na_2_HPO_4_/ Na_2_HPO_4_, pH 7.5 containing 4% PFA at 4°C. delivered with a peristaltic pump at 20 ml/min for 5 min. Brains were then extracted, post-fixed overnight in 4% PFA, and stored at 4°C. 30 μm-thick coronal sections were performed with a vibratome (Leica, Nussloch, Germany) and kept at −20°C in a cryoprotective solution containing 30% ethylene glycol (v/v), 30% glycerol (v/v) and 0.1 M PBS.

### Immunohistochemistry

On day 1, free-floating sections were rinsed three times for 5 min in Tris-buffered saline (TBS, 0.9% NaCl, 0.1 M Tris base, pH 7.5). Sections were then incubated in blocking solution containing 3% normal goat serum, 0.2% triton X-100 (Sigma, Aldrich) and 50 mM NaF for 2 h at room temperature (RT) before overnight 4°C incubation with primary antibodies (see antibody table) diluted in the blocking buffer. On day 2, after three 10-min rinses in TBS, sections were incubated for 90 min with secondary antibodies (see antibody table). The anti-FosB antibody recognizes full-length FosB as well as ΔFosB but at the time point studied (24h post cocaine) all FosB-immunoreactive protein represents ΔFosB (*57*). After three 5-min rinses in TBS, sections were incubated for 5 min with Hoechst (Invitrogen) for nuclei counterstaining. Three 5-min TBS and two Tris buffer (0.1 M Tris base, pH 7.5) washes were performed before sections were mounted in Prolong Gold (Invitrogen).

### Immunoblotting

Mice were killed by decapitation and their heads were immediately snap frozen in liquid nitrogen. Microdiscs of NAc were punched out using disposable biopsy punches (1 mm diameter) (Kai medical) and stored in individual tubes at −80°C. Microdiscs were then homogenized by sonication in a lysis solution containing 50 mM Tris Hcl, 2% SDS, 0.5 M urea diluted in water. Protein concentrations were determined using the BCA assay kit (Pierce, Rockford, IL). A fixed amount of protein (30 μg per lane) was separated by SDS-polyacrylamide gel electrophoresis (12%) before electrophoretic transfer onto nitrocellulose membranes. Membranes were incubated 1 h at RT in a blocking solution containing TBS with 0.1% tween and 5% non-fat dry milk. Membranes were incubated at 4°C overnight with primary antibodies (see antibody table). Membranes were then washed three times in TBS-Tween and then incubated for 1h at RT with secondary antibodies (see antibody table) coupled to horseradish peroxidase (HRP). Immunoreactive bands were detected by chemoluminescent detection (ECL kit, GE Healthcare) and images were acquired using the ImageQuant LAS 4000 (GE Healthcare Life Science). The densitometry of immunoreactive bands was quantified using ImageJ and normalized to the loading control.

### Human brain samples

Brain samples from individuals with a history of substance dependence, with toxicological evidence of current psychostimulant use, and from matched healthy controls (n = 13/group; Table S1) were provided by the Suicide section of the Douglas-Bell Canada Brain Bank (DBCBB). Brains were donated to the DBCBB by familial consent through the Quebec Coroner’s Office, which ascertained the cause of death. Two months after death, psychological autopsies with next-of-kin were conducted, as previously described (*58*).

Dissections of ventral striatum were performed with the guidance of a human brain atlas (*59*) on 0.5 cm-thick formalin-fixed coronal brain sections at an anatomical level equivalent to plate 15 of this atlas (−7.5 mm from the center of the anterior commissure). Tissue was extracted rostral to the anterior commissure and ventral to the tip of the anterior limb of internal capsule and fixed by immersion in formalin until paraffin embedding. The latter was completed using a Leica ASP200S automated processor. Tissue blocks were dehydrated in increasing gradients of alcohol (70%, 95%, 3×100%) for 1.5 hr each, followed by clearing in 3 changes of 100% xylene (2 hr, 2×1.5 hr). The samples were then infiltrated in 3 changes molten paraffin, 3hrs each before embedding. Slices of 6 μm thickness when then prepared with microtome at the histology facility of the ICM institute (Paris).

### Proximity Ligation Assay

Proximity ligation assay (PLA) on mouse brain sections (30 μm-thick) was performed in 48-well plates according to the manufacturer’s instructions for free-floating sections. PLA on human post-mortem paraffin-embedded tissue was performed on sections mounted on superfrost plus slides (Thermo Scientific). 6 μm-thick human caudate-putamen sections were deparaffinized in xylene, rehydrated in graded ethanol series and washed briefly in TBS. For antigen retrieval, sections were boiled for 6 min in sodium citrate buffer (10 mM, pH 6). For brightfield PLA, sections were incubated for 30 min at RT in TBS containing 1% H2O2 to block endogenous peroxidase. After three 5 min rinses with TBS containing 0.1% triton X-100 (TBS-T), sections were incubated 1h at RT with blocking buffer (Duolink blocking buffer for PLA) then with primary antibodies (see antibody table) diluted in the Duolink antibody dilution buffer overnight at 4°C. Anti-rat PLA plus and minus probes were made using the PLA probemaker kit (Sigma Aldrich) with a Goat anti-rat IgG antibody (Jackson Immunoresearch) according to manufacturer’s instructions. Immunofluorescent PLA and the remaining procedures for brightfield PLA were performed as previously described (*31*). For brightfield PLA nuclei were counterstained using the duolink nuclear stain. In this study, we used both single-recognition PLA, using only one primary antibody to detect single antigen and dual-recognition to detect DAR-NMDAR complexes using two primary antibodies.

### Antibodies

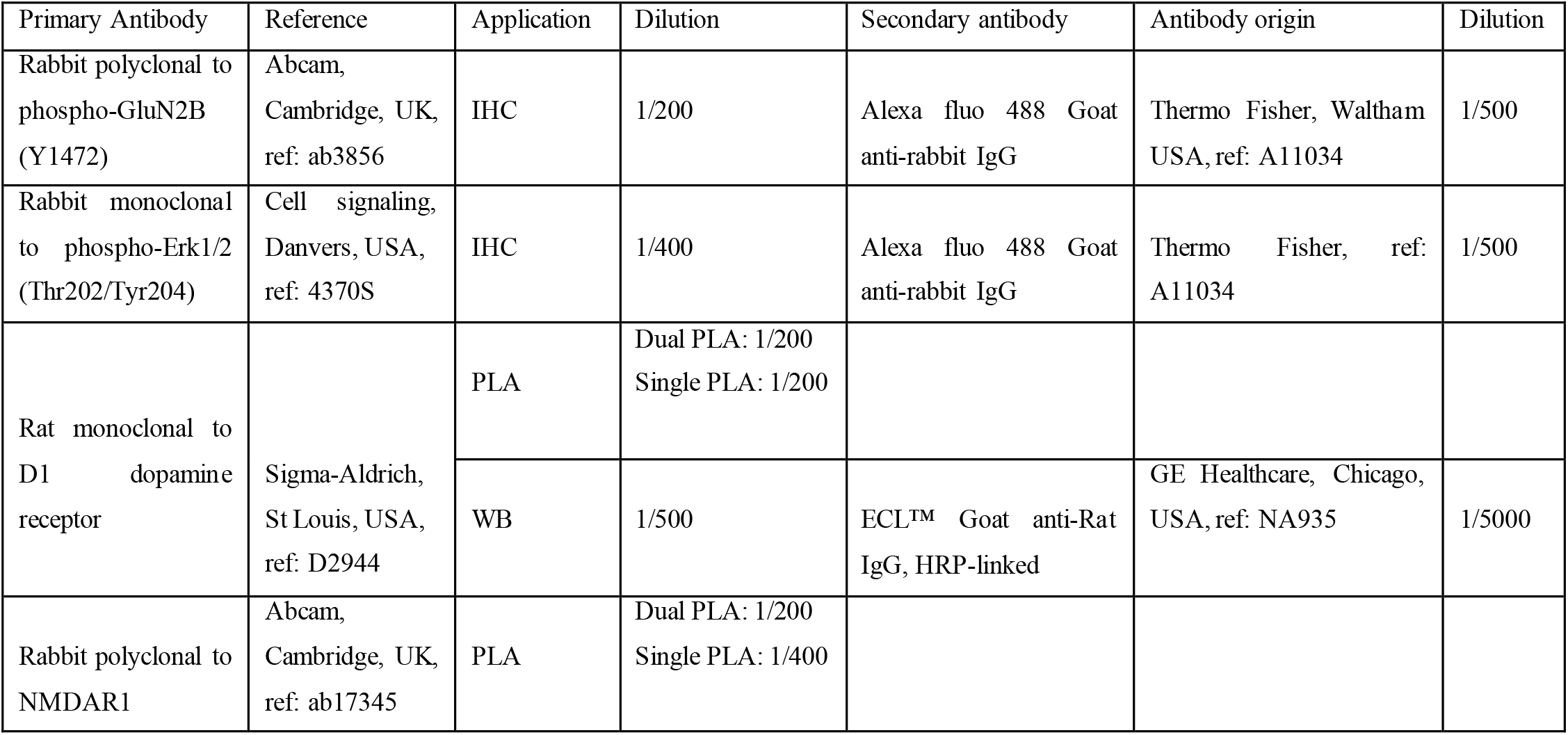
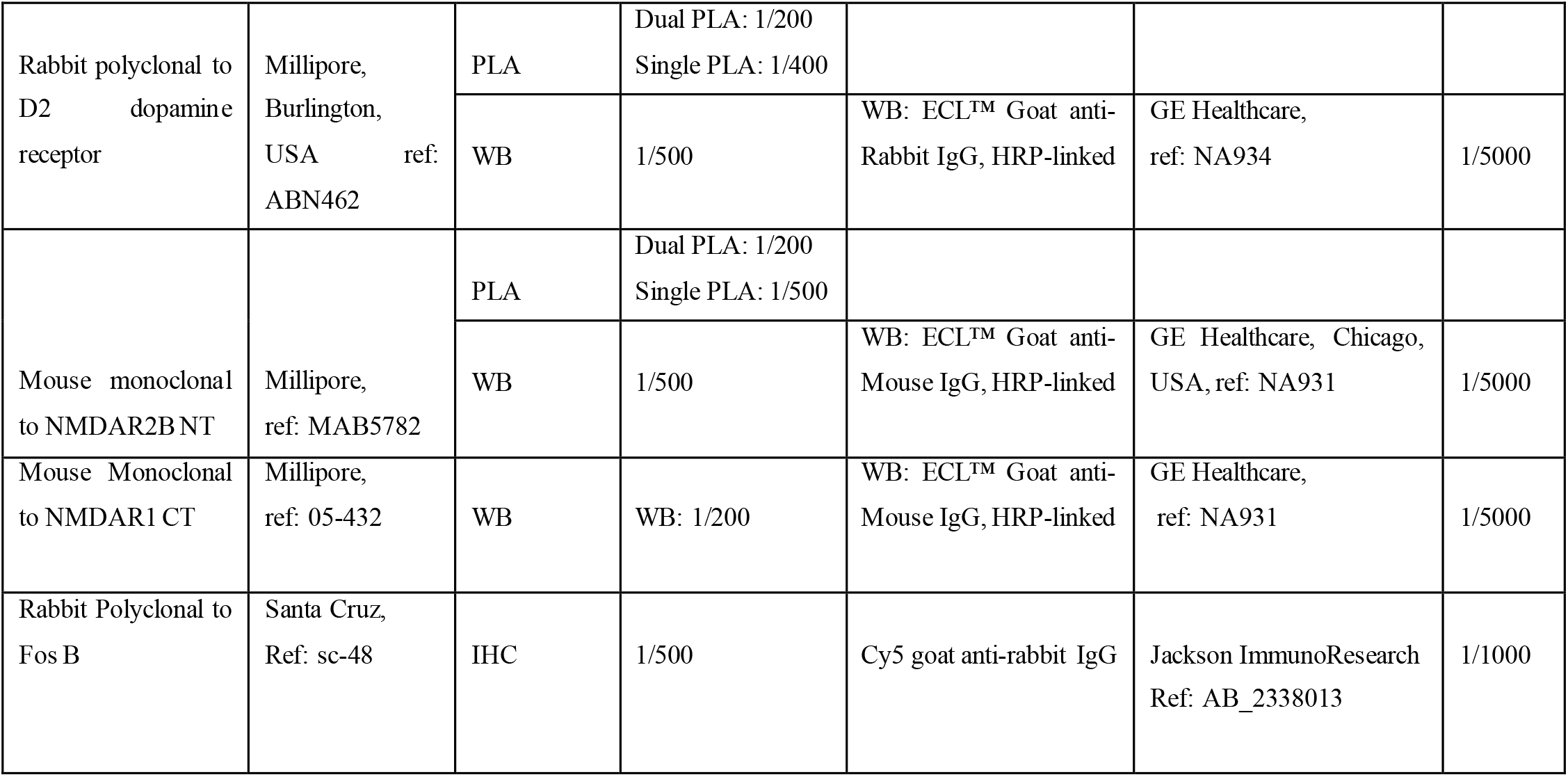

### Image acquisition and analysis

For immunochemistry and fluorescent PLA staining, images were taken with a confocal laser scanning microscope (SP5, Leica) using a 20X and a 63X objective (oil immersion, Leica) respectively. The pinhole was set to 1 Airy unit, excitation wavelength and emission range were 495 and 500-550 nm for green PLA signal, 488 and 500-550 nm for Alexa fluo 488 and 590-650 nm for RFP. Laser intensity and detector gain were constant for all image acquisitions. Images were acquired in a range of 5 μm with a z-step of 0.2 μm. Conditions were run in duplicate and quantifications were made from at least 4 images per condition. Four mice were used for each condition. Maximum projection images were analyzed using ImageJ (National Institutes of Health, Bethesda, MD).

For PLA images quantification, ImageJ was used to construct a mask from RFP positive cells. The mask was then fused with the PLA signal-containing channel using the image multiply function. The PLA punctate signal was quantified on the resulting image in ICY using the spot detector function (detector = scale 3: 65; filtering = Min size: 6; Max size: 30); these parameters were chosen manually from random images to obtain optimal signal-to-noise ratio and minimal false positive in images from negative control conditions. RFP-positive cells showing at least one PLA puncta for one or the other heteromer subtypes were included as this allowed us to identify the MSN subtype analyzed (D1R-MSN v.s D2R-MSN). RFP positive cells which did not present any D1R-GluN1 or D2R-GluN2B PLA signal were discarded.

Mouse brightfield PLA images were taken using a microscope (Leica, DM4000) with a 63X objective. A minimum of 6 random fields per structure per mice were taken.

For human brightfield PLA, whole-slide images were taken using a Zeiss, Axioscan with a 40X magnification and z-stack with a 1 μm step interval. Whole-slide scanned images were visualized with ZEN Blue edition lite software (Zeiss, version 3.1). 25 region of interest (ROI) of 500 μm x 500 μm were generated randomly in the NAc and were exported as TIFF files using the image export plugin. The z-stack with the maximum number of PLA signal in the focus was chosen for each image.

Human and mouse brightfield PLA Images were analyzed using an ImageJ homemade macro that use the Find Maxima tool to detect the PLA punctate with prominence set at >40. Results are represented as the mean PLA signal density per field of views.

For quantification of immunohistochemistry experiments, immunoreactive cells, were analyzed using ImageJ considering the cells with immunofluorescence above a fixed threshold.

### Neuronal survival

Neuronal survival was quantified manually based on Hoescht-counterstained nuclei. Survival was defined as the percentage of viable neurons exhibiting large, uniform nuclei and even distribution of Hoescht among RFP positive cells over RFP positive cells classified as apoptotic based on, at least, two of the following criteria: condensed nuclei, single chromatin clump, small nuclei size, non-circular nuclei shape and increased Hoescht intensity.

### Spine density analysis

Twenty-four hours after the CPP paradigm, mice were perfused and brains were sliced as described above. Dendrite spine analysis was performed on RFP-positive dendrites from D1R-MSN and D2R-MSN identified based on the presence of GFP driven by the co-injection of AAV-PPTA-cre/AAV-pCAG-DIO-eGFP or AAV-PPE-cre/pCAG-DIO-eGFP, respectively. Image stacks were taken using a confocal laser scanning microscope (SP5, Leica). For analysis of D1R-MSN, images were collected through 63x objective with pixel size of 65 nm and z-step of 200 nm. For D2R-MSN, images were collected through 40x objective with pixel size of 95 nm and z-step of 300 nm. The excitation wavelength was 488 for GFP and 561 for RFP, with emission range 500-550 nm and 570–650 nm, respectively. Images were acquired in sequential mode with a Hybrid detector (HyD, Leica). Deconvolution with experimental point spread function from fluorescent beads using a maximum likelihood estimation algorithm was performed with Huygens software (Scientific Volume Imaging). Neuronstudio software was used to reconstruct the dendrite and detect dendritic spines with manual correction.

### Patch clamp recordings

Mice were anesthetized (Ketamine 150 mg/kg / Xylazine 10 mg/kg) and transcardially perfused with aCSF for slice preparation. Coronal 250 μm slices containing the nucleus accumbens were obtained in bubbled ice-cold 95% O_2_/5% CO_2_ aCSF containing (in mM): KCl 2.5, NaH_2_PO_4_ 1.25, MgSO_4_ 10, CaCl_2_ 0.5, glucose 11, sucrose 234, NaHCO_3_ 26, using a HM650V vibratome (Microm, France). Slices were then incubated in aCSF containing (in mM): NaCl 119, KCl 2.5, NaH_2_PO_4_ 1.25, MgSO_4_ 1.3, CaCl_2_ 2.5, NaHCO_3_ 26, glucose 11, at 37°C for 1 h, and then kept at room temperature. Slices were transferred and kept at 31°C in a recording chamber superfused with 2 ml/min aCSF in the continuous presence of 50 μM picrotoxin (Sigma-Aldrich, France, dissolved in DMSO) to block GABAergic transmission. Neurons were visualized by combined epifluorescent and infrared/differential interference contrast visualization using an Olympus upright microscope holding 5x and 40x objectives. Whole-cell voltage-clamp recording techniques were used to measure synaptic responses using a Multiclamp 700B (Molecular Devices, Sunnyvale, CA). Signals were collected and stored using a Digidata 1440A converter and pCLAMP 10.2 software (Molecular Devices, CA). AMPA-R/NMDA-R ratio was assessed using an internal solution containing (in mM) 130 CsCl, 4 NaCl, 2 MgCl_2_, 1.1 EGTA, 5 HEPES, 2 Na_2_ATP, 5 sodium creatine phosphate, 0.6 Na_3_GTP and 0.1 spermine. Synaptic currents were evoked by stimuli (60 μs) at 0.1 Hz through a glass pipette placed 200 μm from the patched neurons. Evoked-EPSCs were obtained at V=+40mV in the absence and presence of the AMPA-R antagonist DNQX. In all cases, 30 consecutive EPSCs were averaged and offline analyses was performed using Clampfit 10.2 (Axon Instruments, USA) and Prism (Graphpad, USA). Pharmacologically isolated EPSC NMDAR decay time, recorded from cells voltage clamped at +40 mV, was fitted with a double exponential function, using Clampfit software, to calculate both slow and fast decay time constants, τfast and τslow, respectively. The weighted time constant (τweighted) was calculated using the relative contribution from each of these components, applying the formula: τw = [(af.τf) + (as.τs)]/(af + as), where af and as are the relative amplitudes of the two exponential components, and τf and τs are the corresponding time constants.

### Cyclic AMP accumulation assay

HEK 293 cells stably expressing the D2R (*60*) were grown on polylysine (Poly-D-lysine hydrobromide, Sigma) and transfected with Tet-On plasmids encoding either the D2R-IL3 or D2R-IL3-scr peptides and the Tag RFP (4 μg per well), complexed with Lipofectamine^®^ 2000 transfection reagent (Invitrogen™), in DMEM (DMEM high glucose GlutaMAX™ Supplement pyruvate, Gibco), and incubated for 5 h. Post-transfection, cells were rinsed and wells filled with DMEM supplemented with 10% FBS and antibiotics to induce D2R expression (Hygromycine 2 μl/ml (Sigma Aldrich); Blasticidine 1,5 μl/ml (Cayla Invivogen ant-bl-1); Tetracycline (Sigma T7660-5g) 1 μl/ml) and incubated overnight. Peptide expression was induced by addition of 1 μg/ml dox solution, incubating for 48 hours. Following the dox treatment, cells were rinsed in DMEM before pretreatment with 1mM IsoButylMethylXanthine (IBMX, Sigma Aldrich) for 15 min. Cells were then stimulated for 30 min with the indicated concentrations of agonist Quinpirole (Tocris), in the presence of 1 mM IBMX and 10 μM Forskolin (Tocris). Endogenous phosphodiesterase activity was stopped by aspiration of the medium and the addition of 0.1 M HCL (300 μl/ well). After centrifugation at 600 g during 10 min, protein concentration of supernatants was quantified by BCA (Uptima, Interchim). Cyclic AMP levels were determined in samples containing 10 μg of protein. The accumulation of cAMP was measured by using a cAMP Enzyme Immunoassay kit (Sigma Aldrich) as described by the manufacturer using Victor3 (Perkin Elmer) plate reader. The curve fit was obtained by GraphPad Prism 8 (GraphPad Software, Inc.).

### Statistical analysis

Results were analyzed with Graphpad Prism (version 8.0.1). Sample size was predetermined on the basis of published studies, pilot experiments and in-house expertise. All data are displayed as mean +/− SEM. Two-tailed Student’s test was used for the comparison of two independent groups. For more than two groups comparison, one-way, two-way or three-way repeated-measures ANOVA were performed followed by Bonferoni Post-hoc test. Data distribution was assumed to be normal and variances were assumed to be homogenous. The main effect and post-hoc statistical significances are given in the appropriate figure legend for each experiment.

## Data availability

The datasets that support the findings of this study are available from the corresponding author upon reasonable request.

## Acknowledgments

This work was supported by the Centre National de la Recherche Scientifique (CNRS); the Institut National de la Santé et de le Recherche Médicale National (INSERM); Sorbonne Université, Faculté des Sciences et Ingénierie; Université de Bordeaux; Institut National de Recherche pour l’Agriculture, l’Alimentation et l’Environnement (INRAE); Université Côte d’Azur; the Agence Nationale pour la Recherche (ANR; ANR-15-CE16-001 to P.V and P.T; ANR-18-CE37-0003-02 to J.B and P.V; ANR-10-IDEX-03-02 and ANR-16-CE16-0022 to P.T); the Fondation pour la Recherche Médicale (FRM; DPA20140629798 to J.C; DEQ20180339159 to J.B.); Institut de Recherche en Santé publique (IReSP) Aviesan APP-addiction 2019 (to P.V, P.T and J.B); NARSAD Young Investigator Grants from the Brain and Behavior Foundation (to P.T); the BioPsy labex excellence cluster (to J.C and P.V); the labex BRAIN (to P.T) and NIH grant MH54137 (to J.A.J.). The Douglas-Bell Canada Brain Bank is funded by platform support grants from the RQSHA and HBHL (CFREF). A.A and R.W. are recipients of PhD fellowship from the French Ministry of Research; A.P is the recipient of a PhD fellowship from the “Ecole Universitaire de Recherche” (*EUR Neuro,* Bordeaux Neurocampus); E.S.J is the recipient of a fourth-year PhD fellowship from the FRM. Authors would like to thank the imaging facility of the IBPS and the histology facility of the Institut du Cerveau et de Moëlle épinière (ICM).

## Authors contributions

A.A performed most PLA, viral injections, behavioral studies, immunohistochemistry, confocal imaging, biochemistry and statistical analysis with the help of E.S.J; M.C.A; V.K; S.B, V.O and R.W. Dendritic spines analysis was performed by N.H and A.A. Electrophysiological recordings were performed by P.P and S.P.F. Viruses were designed by P.V and A.P.B and produced by C.J and A.P.B and V.S.P and A.P performed cAMP assays. G.T and N.M obtained, characterized and provided human brain samples. Y.Z and J.A.J. helped A.A for PLA experiments from human tissues and related quantifications. A.A; J.C; P.T; J.B and P.V designed experiments and wrote the manuscript, which was edited by the other authors.

## Competing interests

The authors declare no competing interests

## Notes

### Competing Interest Statement

The authors have declared no competing interest.

