## Supplementary Materials for "Disrupting D2-NMDA receptor heteromerization blocks the rewarding effects of cocaine but preserves natural reward processing"

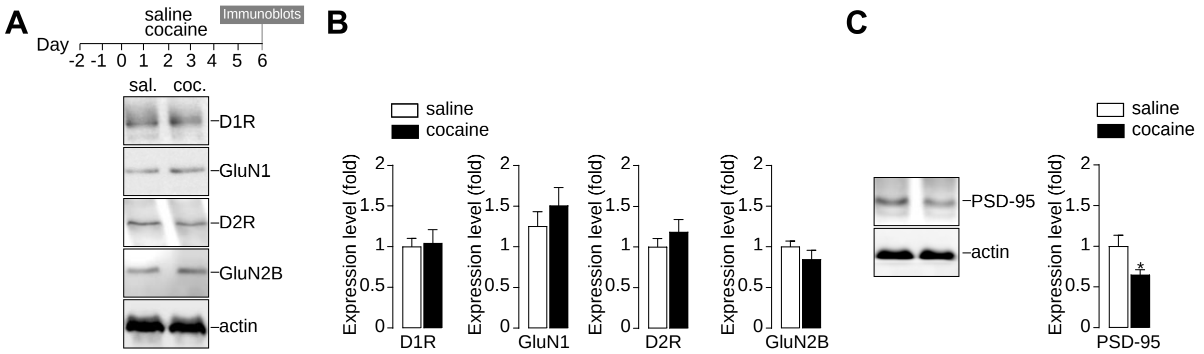


**Fig. S1. Repeated cocaine exposure does not alter expression levels of D1R, D2R and GluN1/2B subunits of NMDAR in mice.** (**A**) Top: Experimental time frame of saline or cocaine (15mg/kg) treatments. Bottom: representative D1R, GluN1, D2R, GluN2B and actin immunoblots performed 24h after the last saline (sal.) or cocaine (coc.) injection. (**B**) Quantifications of D1R expression levels: Two-sided Student's t-test, t = 0.2263 df. = 21, P =0.8232 n=11-12 mice/group; GluN1 expression levels: Two-sided Student's t-test, t = 1.001 df. = 21, P =0.3282 n=11-12 mice/group; D2R expression levels: Two-sided Student's t-test, t = 0.992 df. = 20, P =0.33326 n=11 mice/group and GluN2B expression levels: Two-sided Student's t-test, t = 1.163 df. = 20, P =0.2587 n=11 mice/group normalized to actin levels and represented as fold relative to the saline-treated group. (**C**) Representative PSD-95 and actin immunoblots and quantifications of PSD-95 expression levels (normalized to actin) relative to the saline-treated group. Two-sided Student's t-test, t = 2.272 df. = 21, *P =0.0338 n=11-12 mice/group. Error bars denote s.e.m.

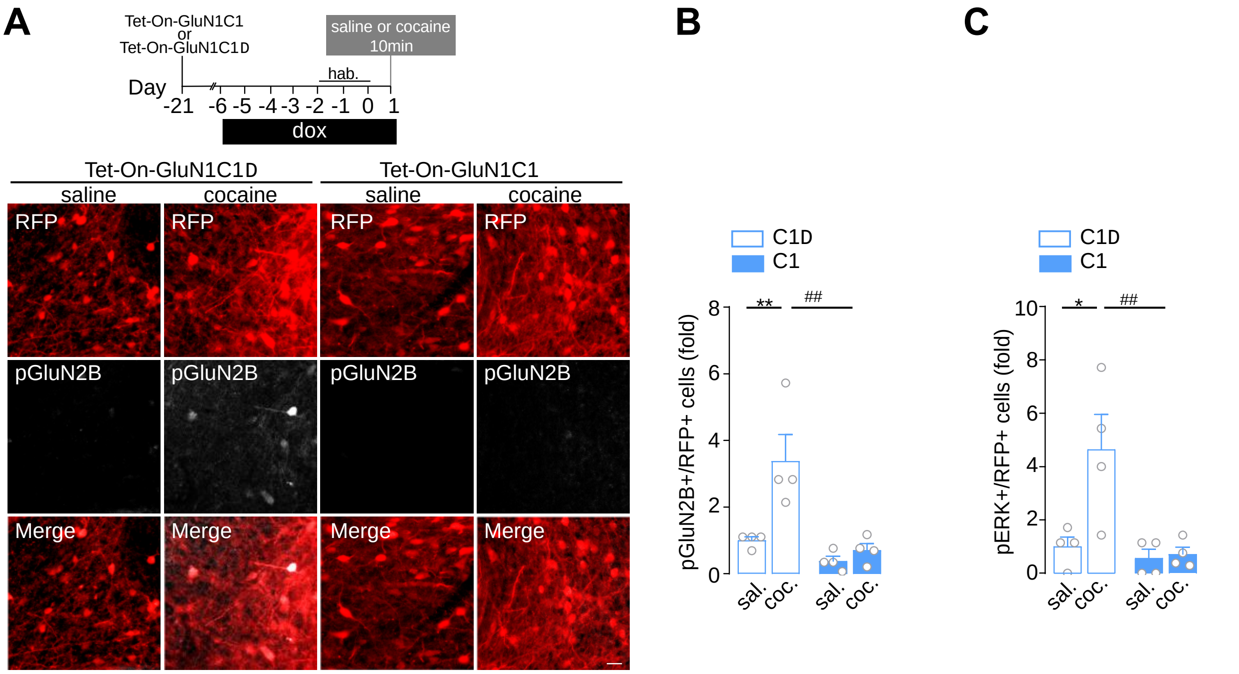


**Fig. S2. The inhibition of of D1R-GluN1 heteromerization in the NAc with AAV Tet-On-GluN1C1 blocks cocaine-mediated GluN2B phosphorylation and ERK activation**. (**A**) Top: Time frame of AAV-Tet-On injection, doxycycline (dox) treatment and saline or cocaine (15 mg/kg) administration. Mice were perfused 10 min after a single injection of saline or cocaine. Bottom: Representative images of RFP expression and GluN2B phosphorylation (pGluN2B) in saline or cocaine-treated mice infected with Tet-On-GluN1C1Δ or Tet-On GluN1C1. Scale bar: 30 μm. (**B**) Quantifications of pGluN2B in RFP positive neurons infected with the Tet-On-GluN1C1Δ (C1Δ) or Tet-On-GluN1C1 (C1) AAVs represented as fold relative to the saline-treated group infected with C1Δ. Two-way ANOVA: virus effect, $F(1, 12) = 10.36$, **P =0.0083, saline C1Δ vs cocaine C1Δ; ##P=0.0035, cocaine C1 vs cocaine C1Δ, n=4 mice/group. (**C**) Quantifications of ERK phosphorylation (pERK) in RFP neurons infected with the Tet-On-GluN1C1Δ (C1Δ) or Tet-On-GluN1C1 (C1) AAVs represented as fold relative to the saline-treated group infected with C1Δ. Two-way ANOVA: virus effect, $F(1, 12) = 7.031$, *P=0.0164, saline C1Δ vs cocaine C1Δ; ##P=0.010, cocaine C1 vs cocaine C1Δ, n=4 mice/group. Error bars denote s.e.m.

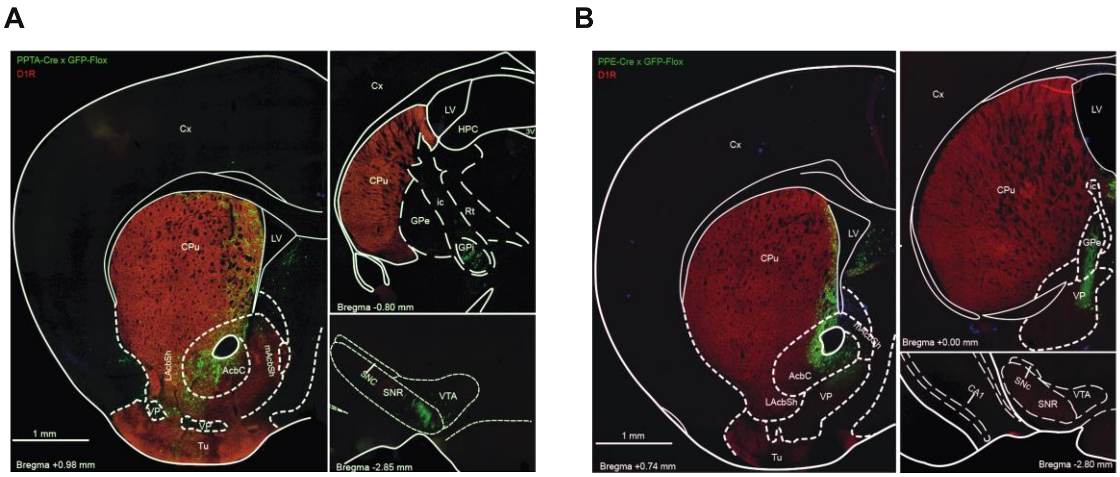


**Fig. S3. Virally-mediated tagging of D1R- and D2R-MSN**. (**A**) Illustrative coronal sections obtained from mice injected with the mixture of AAV-PPTA-Cre and AAV-DIO-GFP in the NAc and the dorso-medial striatum to tag D1R-MSN. Note that, this combination of AAV yielded, as expected, a GFP signal in projections structures of D1R-MSN, e.g. the GPi, the SNR and the VTA, as expected for D1R-MSN. (**B**) Mice injected with the mixture of AAV-PPE-Cre and AAV-DIO-GFP in the NAc and the dorsal striatum to tag D2R-MSN. This combination of AAV leads to GFP staining in the GPe and VP, but in the SNR and the VTA. A D1R immunolabeling (red) was performed to facilitate the visualization of distinct brain regions. Abbreviations: 3V: 3rd ventricle; AcbC: accumbens nucleus core; Cpu: Caudate putamen; Cx: cortex; GPe: external globus pallidus; GPi: internal globus pallidus HPC: hippocampus; ic: internal capsule; LAcbSh: lateral accumbens shell; LV: lateral ventricle; mAcbSh: medial accumbens shell; Tu: olfactory tubercle; VP: ventral pallidum; Rt: reticular nucleus; SNR: substantia nigra, pars reticulata; SNC: substantia nigra, pars compacta; VTA: ventral tegmental area

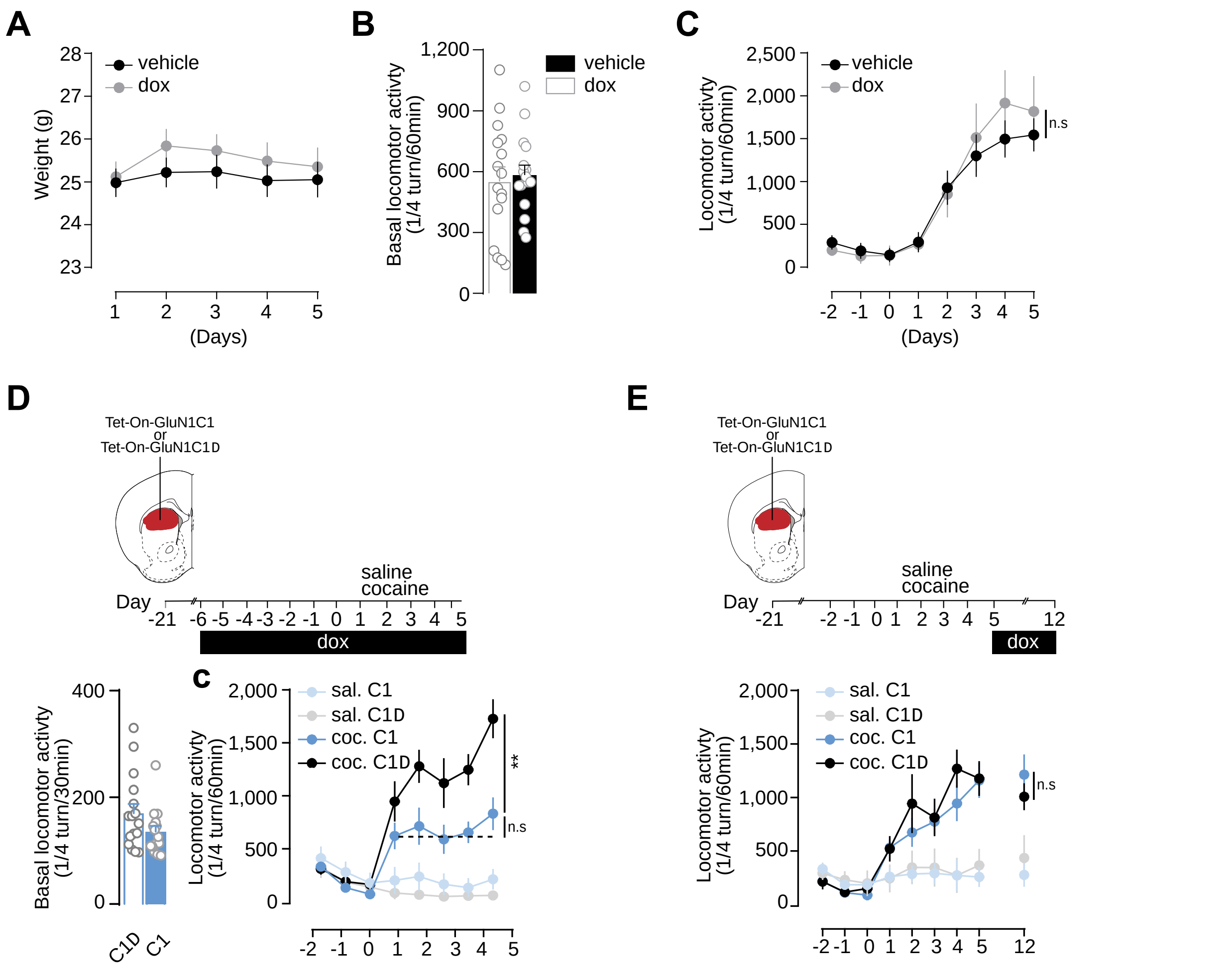


**Fig.S4. Innocuity of doxycycline treatment in mice that are not injected with an AAV-Tet-On and impairment of the development, but not maintenance, of cocaine-induced locomotor sensitization upon inhibition of D1R-GluN1 heteromerization in the dorsal striatum.** (**A**) Mice were treated for 7d with doxycycline (dox) prior to and during body weight measurement. Two-way ANOVA: doxycycline effect, $F(1, 30) = 0.6105$, $P > 0.999$, dox vs vehicle, n=16 mice/group. (**B**) Lack of effect of dox administration on basal locomotion. Two-sided Student's t-test, $t=0.356$ $df.=30$, $P=0.723$ n=16 mice/group. (**C**) Cocaine-induced locomotor sensitization of mice treated or not with dox prior and during cocaine injections. Two-way ANOVA: doxycycline effect, $F(1, 13) = 0.152$, n. $P > 0.999$, cocaine dox vs cocaine vehicle locomotion on d5, n=7-8 mice/group. (**D**) Experimental time frame to study the consequences of inhibiting D1R-GluN1 heteromerization in the dorsal striatum on basal locomotion and development of cocaine-induced locomotor sensitization. Basal locomotor activity: Two-sided Student's t-test, t = 1.653 df. = 29, P =0.1092, n=15-16 mice/group. Cocaine-induced locomotor sensitization: Three-way ANOVA: virus effect, F (1, 224) = 9.643, **P<0.001, n.s P>0.999, n=6-10 mice/group. (**E**) Experimental design to study the role of D1R-GluN1 heteromerization in the dorsal striatum on the maintenance of cocaine-induced locomotor sensitization. Locomotor activity in each group: Three-way ANOVA: virus effect, F (1, 243) = 0.2522, n.s P >0.999, n=7-8 mice/group. n.s not significant. Error bars denote s.e.m.

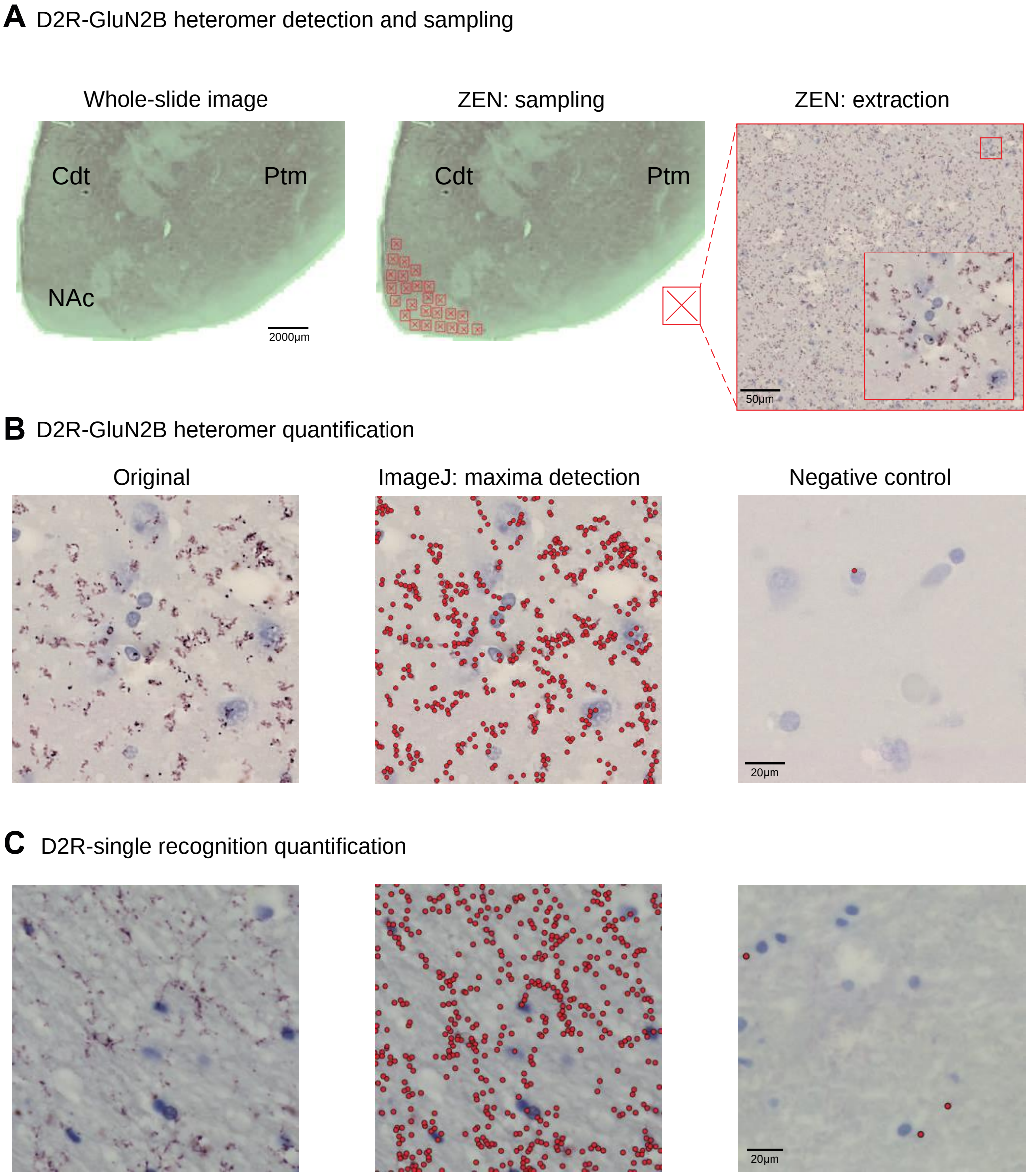

**Fig.S5. Analysis of the PLA signal from human post-mortem caudate putamen samples**. (**A**) From a whole-slide image of human caudate putamen slice (left), the ZEN software allows the extraction of 25 high magnification images randomly selected in a region of interest corresponding to the nucleus accumbens (middle). Extracted images are used to automatically detect the PLA signal (right). (**B**) Example image of the D2R-GluN2B PLA signal (left) and its detection by a custom-made macro for local maxima detection (middle panel). Right panel: example image showing the absence of PLA signal in the negative control where the GluN2B antibody is omitted during the PLA. (**C**) Same as B in the case of D2R single recognition. Abbreviations: Cdt: caudate; NAc: nucleus accubens; Ptm: putamen;

| Control samples | | | | | | |
|---|---|---|---|---|---|---|
| Gender | Age | Cause of death | PMI | pH value | Substances at death | Axis 1 dependence |
| Male | 28 | suicide | 16 | 6,93 | Nil | Nil |
| Male | 54 | suicide | 36 | N/A | Nil | Nil |
| Male | 26 | suicide | 55 | 7,30 | Nil | Nil |
| Male | 54 | suicide | 19 | 7,10 | Nil | Nil |
| Male | 29 | suicide | 56,5 | 6,40 | Nil | Nil |
| Male | 72 | Natural | 51 | N/A | N/A | Nil |
| Male | 18 | Natural | 68.5 | 6,87 | Nil | Nil |
| Female | 27 | Accident | 79,5 | 5,98 | N/A | Nil |
| Male | 31 | Natural | 101 | 6,80 | Nil | Nil |
| Male | 71 | Natural | 17 | 6,20 | Nil | Nil |
| Male | 81 | Accident | 26,75 | 5,80 | N/A | Nil |
| Male | 55 | Natural | 21 | 6,70 | Nil | Nil |
| Female | 40 | Natural | 106,5 | 6,50 | N/A | Nil |
| **Samples from addict individuals** | | | | | | |
| Gender | Age | Cause of death | PMI | pH value | Substances at death | Axis 1 dependence |
| Male | 39 | Suicide | 36,75 | 6,74 | **Cocaine and metabolites** | Substance dependence (alcohol) |
| Female | 51 | Suicide | 33,75 | N/A | Ethanol, **cocaine and metabolites** | Substance dependence |
| Male | 39 | Suicide | 66,75 | 6,70 | **Cocaine and metabolites** | Substance dependence (**cocaine**) |
| Male | 38 | Suicide | 67 | 6,50 | **Cocaine and metabolites**, antidepressants (SNRI), benzodiazepines, cannabinoids and metabolites | Substance dependence |
| Male | 53 | Suicide | 65,75 | 6,50 | **Cocaine and metabolites**, ethanol, lidocaine | Substance dependence |
| Male | 52 | Suicide | 86,5 | 6,20 | **Cocaine**, ethanol | Substance dependence |
| Male | 37 | Suicide | 18,5 | 6,90 | **Cocaine and metabolites**, antidepressants (SSRI) | Substance dependence |
| Male | 45 | Suicide | 38 | 6,50 | **Cocaine and metabolites**, cannabinoids and metabolites | Substance dependence |
| Male | 24 | Accident | 77,25 | 6,33 | **Cocaine and metabolites**, opiates, cannabinoids and metabolites | Substance dependence (**cocaine**) |
| Male | 51 | Accident | 81,75 | 6,80 | Opioids, diphenhydramine, benzodiazepines | Substance dependence (**cocaine**) |
| Male | 32 | Accident | 32,25 | 6,20 | **Amphetamines, methamphetamines**, diphenhydramine | Substance dependence (alcohol) |
| Male | 50 | Natural | 100 | 6,50 | **Cocaine and metabolites** | Substance dependence (**cocaine**) |
| Male | 28 | Accident | 43,5 | 6,50 | **Cocaine and metabolites**, ethanol | Substance dependence |

**Table S1. Detailed human subject information.** PMI: post-mortem interval
